# Genome-scale transcriptional regulatory network models of psychiatric and neurodegenerative disorders

**DOI:** 10.1101/190959

**Authors:** Jocelynn R. Pearl, Dani E. Bergey, Cory C. Funk, Bijoya Basu, Rediet Oshone, Paul Shannon, Leroy Hood, Nathan D. Price, Carlo Colantuoni, Seth A. Ament

## Abstract

Genetic and genomic studies suggest an important role for transcriptional regulatory changes in brain diseases, but roles for specific transcription factors (TFs) remain poorly understood. We integrated human brain-specific DNase I footprinting and TF-gene co-expression to reconstruct a transcriptional regulatory network (TRN) model for the human brain, predicting the brain-specific binding sites and target genes for 741 TFs. We used this model to predict core TFs involved in psychiatric and neurodegenerative diseases. Our results suggest that disease-related transcriptomic and genetic changes converge on small sets of disease-specific regulators, with distinct networks underlying neurodegenerative vs. psychiatric diseases. Core TFs were frequently implicated in a disease through multiple mechanisms, including differential expression of their target genes, disruption of their binding sites by disease-associated SNPs, and associations of the genetic loci encoding these TFs with disease risk. We validated our model’s predictions through systematic comparison to publicly available ChIP-seq and TF perturbation studies and through experimental studies in primary human neural stem cells. Combined genetic and transcriptional evidence supports roles for neuronal and microglia-enriched, MEF2C-regulated networks in Alzheimer’s disease; an oligodendrocyte-enriched, SREBF1-regulated network in schizophrenia; and a neural stem cell and astrocyte-enriched, POU3F2-regulated network in bipolar disorder. We provide our models of brain-specific TF binding sites and target genes as a resource for network analysis of brain diseases.

## Introduction

Convergent evidence suggests that altered transcriptional regulation is a prominent mechanism of common human diseases, including psychiatric and neurodegenerative disorders. Many disease states are accompanied by characteristic, tissue-specific changes in gene expression. Neurodegenerative diseases such as Alzheimer’s disease and Huntington’s disease involve progressive changes in the expression of thousands of genes in vulnerable brain regions^1,2^. Psychiatric disorders such as schizophrenia, bipolar disorder, major depression, and autism also involve brain gene expression changes, with replicable changes in the expression of hundreds of genes observed in neocortical regions that influence cognition and emotional control^3-7^.

While non-genetic factors could contribute to brain gene expression changes in psychiatric and neurodegenerative disorders ‐‐ including effects of medications, lifestyle factors, and differences in cell type distributions ‐‐ multiple lines of evidence point to a substantial genetic contribution. Hundreds of genetic haplotypes associated with risk for psychiatric and neurodegenerative disorders co-localize with gene expression quantitative trait loci (eQTLs)^8,9^. Genetic variants associated with risk for common diseases are enriched in promoters and enhancers^10^, including regions of open chromatin marked by DNase I hypersensitivity^11^. Thus, it has been proposed that the causal variants at many risk loci alter the expression of nearby target genes via mechanisms such as changes in transcription factor binding.

Genetically encoded changes in transcription factors (TFs) and other transcriptional regulatory proteins may also contribute to disease risk. Both common and rare genetic variation associated with psychiatric disorders are enriched for genes involved in transcriptional regulation and chromatin-remodeling pathways^12,13^. 29 TFs are located at genome-wide significant risk loci from large-scale GWAS of schizophrenia or bipolar disorder^14–17^. Additional TFs have been implicated in risk for these diseases due to their disruption by rare variants^18^.

We hypothesized that psychiatric and neurodegenerative disorders involve dysregulation within networks of TFs, TF binding sites, and TF target genes in the brain. Disease related changes in transcriptional networks could occur either at the genetic level (i.e., disease-associated variation in the DNA sequence) or at the transcriptional level (i.e., changes in gene expression), or by both mechanisms. Previous studies have used gene co-expression to identify disease-perturbed networks in the brain^1,7,19,20^ and have observed that a subset of differentially expressed networks are enriched for genes that are also associated with genetic risk for the same disease^1,7^. Other studies have utilized epigenomic profiling and eQTLs to annotate non-coding genetic variation associated with risk for psychiatric and neurodegenerative disorders^4,9^, and to fine-map transcriptional regulatory mechanisms at specific disease risk loci^21^. These studies support the hypothesis that networks of TFs, TF binding sites, and TF target genes to brain diseases. However, genome-wide studies have not previously integrated all of these levels of TRN organization to understand the transcriptional regulatory architecture of brain diseases.

Here, we reconstructed a transcriptional regulatory network (TRN) model that predicts the genomic binding sites and target genes of 741 transcription factors in the human brain by integrating tissue-specific DNase-seq footprinting with TF-gene co-expression. We used our TRN model to predict core TFs that regulate transcriptomic changes in psychiatric and neurodegenerative diseases, as well as disease-associated SNPs that disrupt transcription factor binding sites. We integrate these results to characterize *cis-* and trans-acting mechanisms linking transcriptional regulatory networks to disease (Fig. S1).

## Results

### Reconstruction of a transcriptional regulatory network model for the human brain

To predict the genome-wide binding sites for transcription factors in the human brain, we performed digital genomic footprinting analysis with 15 DNase-seq experiments from human brain regions and cell types generated by the ENCODE project (Table S1). DNase I cleavage patterns predict occupied binding sites for TFs and other DNA binding proteins^22,23^. We downloaded raw sequencing reads from each DNase-seq experiment (www.encodeproject.org), aligned the reads to the human genome with SNAP^24^, identified DNase I hypersensitive regions with F-seq^25^, and located footprints of DNA binding proteins with Wellington^26^. We intersected DNase I footprints with DNA sequence motifs from JASPAR^27^, UniProbe^28^, and SwissRegulon^29^ to predict binding sites for specific TFs, focusing on 741 TFs-approximately half of all human TFs ‐‐ that are expressed in the human brain^30^ and have known sequence specificity. We used the union of TF binding sites predicted across all 15 DNase-seq experiments to identify 2,637,487 DNase hypersensitive sites (DHSs), containing 1,121,670 footprints (Supplementary Dataset 1).

To predict TF-target gene interactions, we integrated our model of brain TFBSs with evidence of co-expression between TF-gene pairs (Fig. 1a). The model was trained using gene expression data from five of the six brains in the Allen Human Brain Atlas^30^, leaving gene expression profiles from the final brain as a test set. Each brain in this dataset was microdissected into ̃350-900 tissue samples, with each sample representing the expression levels of transcripts in ̃10^4^ cells and mapped onto a reference atlas with 862 brain structures^31^. Thus, the Allen Human Brain Atlas defines a cellular resolution map of gene expression in the human brain.

**Figure 1.**
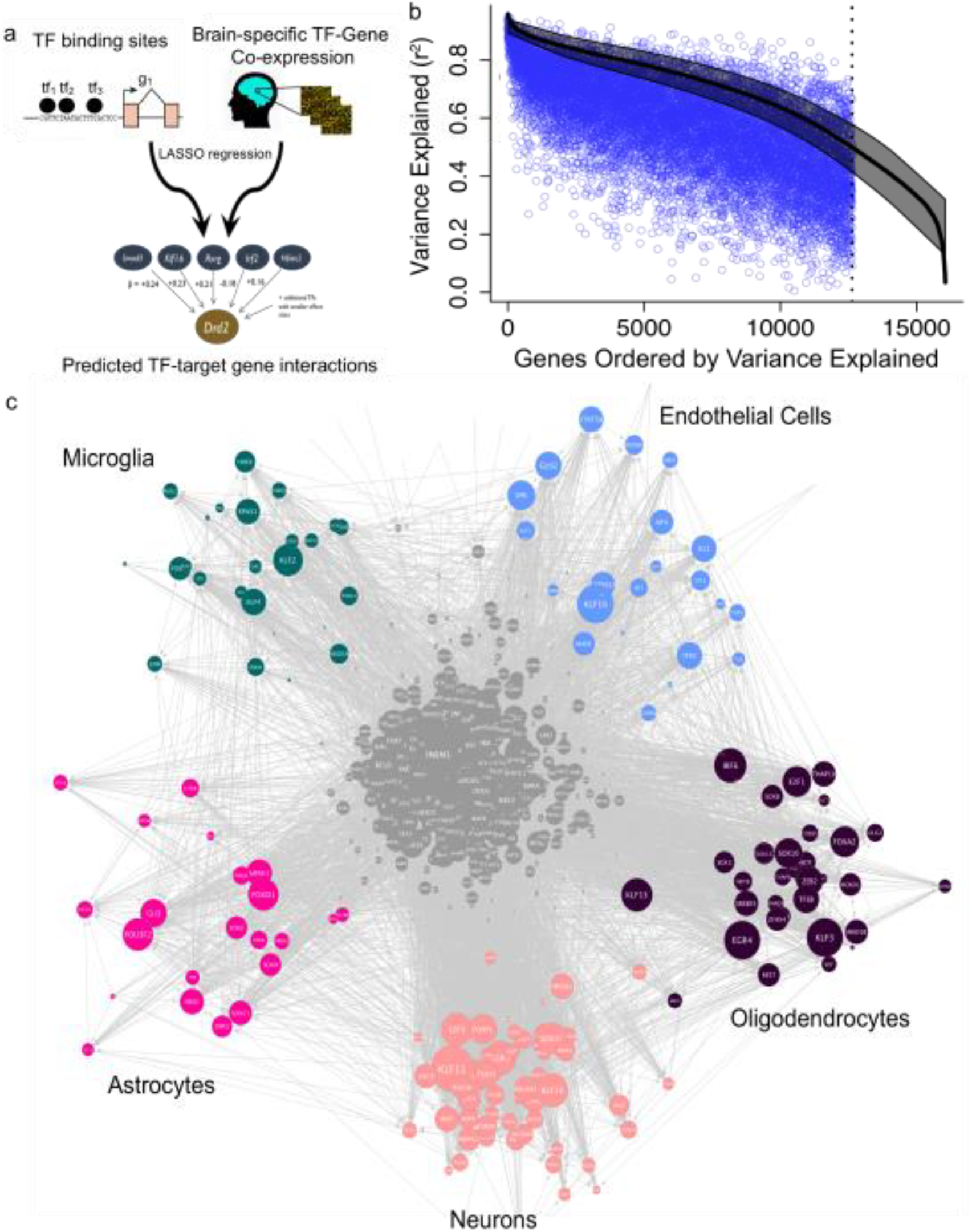
Reconstruction of a transcriptional regulatory network (TRN) model for the human brain. (a) The genomic binding sites and target genes for 741 TFs were predicted by integration of digital genomic footprinting with TF-gene coexpression and LASSO regression. In the LASSO regression model, the expression of each gene is predicted based on the expression levels of transcription factors whose binding sites are enriched +/-10kb from the transcription start site. (b) LASSO regression models for each gene were evaluated by comparing predicted vs. observed expression levels (Pearson’s r^2^) in training and in test sets from the Allen Human Brain Atlas. (c) Predicted regulator interactions among the 741 TFs, and enrichment of each TF’s target genes in one of the major brain cell types (FDR < 0.05). Node size indicates out-degree in the TF-to-TF network.

We applied two algorithms to predict TF-target gene interactions: (i) LASSO regression, in which combinations of TFs are used as predictors of the target gene’s expression^32–35^, and (ii) Pearson correlation between TF-gene pairs. TFBSs were used as constraints, so that only those TFs whose TFBSs were enriched +/-10 kb from a gene’s transcription start site were considered as candidate regulators. We constructed separate models for each of the five brains. We then created a final, consensus model, which retained TF-target gene interactions that were supported by both LASSO and Pearson correlation in at least two of the five brains. In addition, we removed genes for which the LASSO regression model explained <50% of the variance in gene expression. The resulting transcriptional regulatory network (TRN) model includes 741 TFs, 11,093 target genes, and 201,218 TF-target gene interactions (Supplementary Dataset 2).

We systematically evaluated the accuracy of our model’s predictions through comparison to four independent datasets. First, we evaluated co-expression between predicted TF-target gene pairs in 893 Allen Human Brain Atlas samples held out from TRN reconstruction. The expression of 96% of TF-target gene pairs was correlated in this test set (p < 0.05; 75% of TF-target gene pairs strongly correlated at |r| > 0.25). In addition, the prediction accuracy of LASSO regression models learned in training data was strongly correlated to performance in the holdout samples ‐‐ training vs. test set accuracy: r = 0.75; >50% of expression variance explained in the test set for 8,369 of the 11,093 genes in our TRN model; Fig. 1b).

Second, we compared our TRN model to direct targets of 402 of the 741 TFs in our TRN model using ChIP-seq data from diverse human tissues and cell types collected in the GTRD database^36^. 40% of the predicted TF-target gene interactions in our TRN model involving these 402 TFs were supported by a ChIP peak for that TF located <10kb from the TSS of its predicted target gene (more than expected by chance: p << 1e^-4^ by permuting TRN edges; Fig. S2a).

Third, we compared our TRN model to functional target genes of 200 TFs derived from TF knockout, knockdown, or overexpression experiments in various human and mouse tissues and cell types, collected in the CREEDS database^37^. None of these single-gene perturbation studies were performed in human brain or in human neuronal or glial cell lines. Nonetheless, 7% of the predicted TF-target gene interactions in our model involving these 200 TFs were supported by differential expression of the target gene following perturbation of the TF, significantly more than expected by chance (p << 1e-4 by permuting TRN edges; Fig. S2b).

Finally, since many TFs are known to regulate cell-type specific processes in the brain^38^, we asked whether the predicted target genes for each TF were enriched in specific brain cell types and whether these cell-type enrichments were concordant with the expression patterns of the TFs themselves. Using gene expression profiles from purified neurons, astrocytes, oligodendrocytes, microglia, or endothelial cells^39^ and the pSI algorithm^40^, we identified 135 cell-type specific TFs and 147 cell-type enriched TF-target gene modules (Fig. 1c, Table S2). 47 TFs could be assigned to a specific cell-type by both methods. Among these 47 TFs, we observed 91% concordance in cell-type assignments (inter-rater reliability: Cohen’s kappa = 0.89, p << 2e-16). Our model correctly assigned known cell-type markers such as *OLIG2* in oligodendrocytes^41^ and *NEUROD2* and *NEUROD6* in neurons^42,43^. These results suggest that a subset of TRN modules reflect cell-type specific processes. Non-cell-type-specific TFs may regulate a wide variety of biological functions that are active in multiple cell types.

Taken together, the network validation steps described above support the statistical robustness, mechanistic accuracy, and biological interpretability of TF-target gene interactions from our brain TRN model. Since most of the ChIP-seq and single-gene perturbation studies used for validation were performed in non-brain tissues, the number of true positive predictions is likely higher than the estimates from these data.

### Key regulators of disease-related transcriptional changes in prefrontal cortex

Next, we sought to identify TFs that are key regulators of gene expression changes in brain diseases. We focused on gene expression changes in the prefrontal cortex (PFC), a neocortical region that is involved in both emotional control and cognition^44^. Altered prefrontal cortex structure and function are implicated in a wide range of neurodevelopmental, psychiatric, and neurodegenerative disorders^45–50^.

We studied post-mortem prefrontal cortex gene expression profiles from 1,372 human subjects, using data from two to three independent cohorts from each of five common brain diseases: schizophrenia (SCZ)^4,19,51^, bipolar disorder (BD)^51,52^, major depression disorder (MDD)^51,53^, autism spectrum disorder (ASD)^7,54^, and Alzheimer’s disease (AD)^1,55^, as well as non-diseased controls (Table S3). We identified differentially expressed genes in the cases vs. controls from each dataset and tested for over-representation of each TF’s target genes among these differentially expressed genes (Fisher’s exact test). We considered a TF to be a key regulator in a disease if its predicted target genes were over-represented among the differentially expressed genes in that disease, with a meta-analytic q-value < 0.05 across all cohorts and a p-value < 0.05 in at least two independent cohorts. We identified 69 key regulators for schizophrenia, 13 for bipolar disorder, none for major depression, 58 for autism spectrum disorder, and 78 for Alzheimer’s disease (Figure 2b, Tables S4-S7).

**Figure 2.**
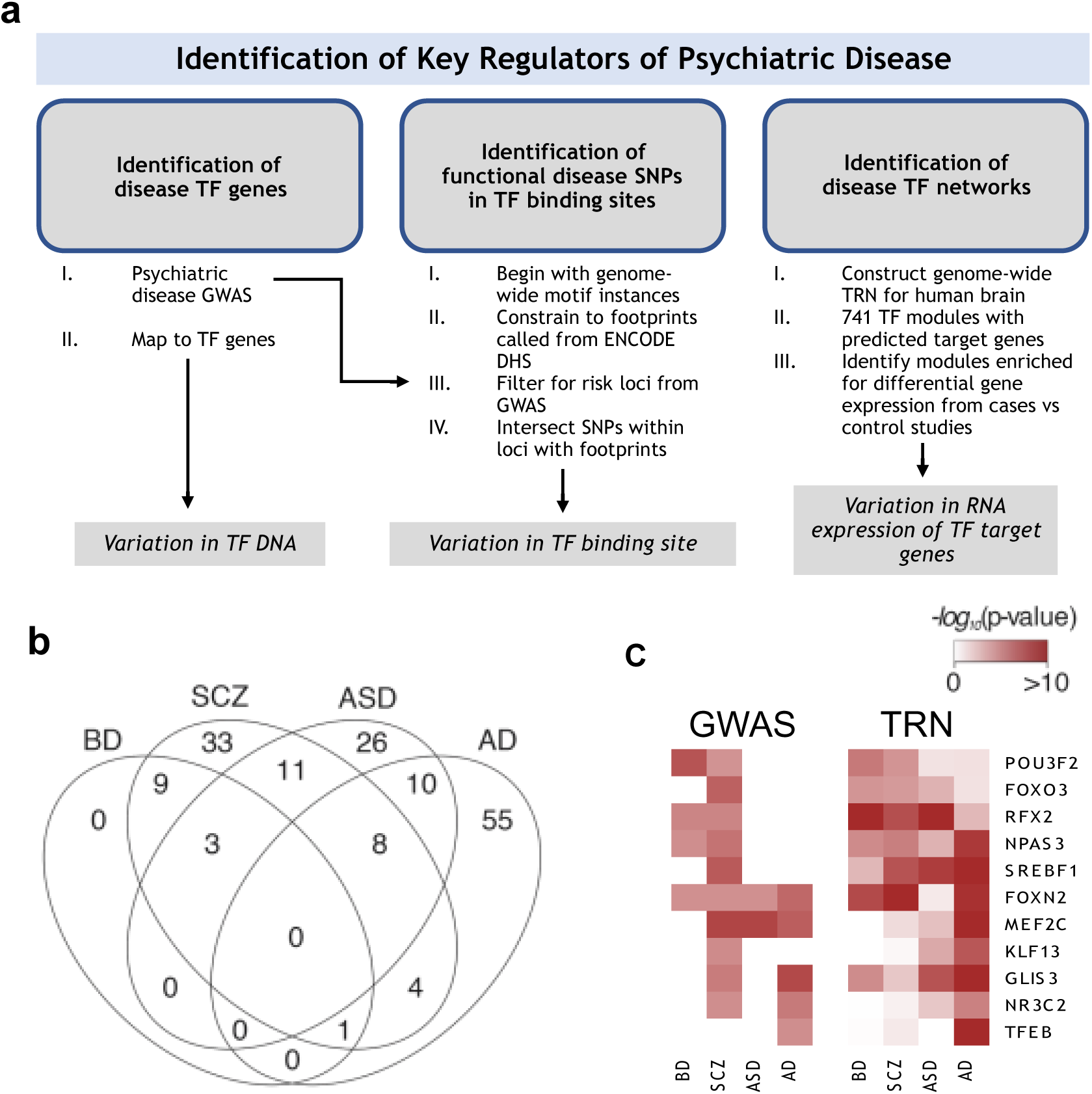
Key regulators of disease-related transcriptional changes in prefrontal cortex. (a) Schematic describing the approach for identifying key regulators in psychiatric disorders (b) Venn diagram describing overlap of key regulators in schizophrenia (SCZ), bipolar disorder (BD), autism spectrum disorder (ASD), and Alzheimer’s disease (AD). (c) ‒log10(p-values) for genetic associations of TF loci and enrichment of TF’s target genes among disease-specific differentially expressed genes for 13 TFs with both a genetic association and a target gene enrichment in one of the four diseases. GWAS p-values are based on studies in the GWAS catalog (Table S8).

We found significant overlap between regulators of disorders with psychotic features (schizophrenia vs. bipolar disorder, 13 shared TFs, 1 expected by chance, p = 1.3e-14; Fig. 2b) and between disorders with a strong neurodevelopmental component (schizophrenia vs. autism, 22 shared TFs, 5 expected by chance, p = 3.1e-9). Intriguingly, while key regulator TFs for schizophrenia were shared with both bipolar disorder and autism spectrum disorder, there was very little overlap between bipolar disorder and autism (3 shared TFs, 1 expected by chance, p = 0.07), suggesting that distinct transcriptional networks underlie the psychotic and neurodevelopment dimensions of schizophrenia. Also, there was relatively little overlap between regulators of bipolar disorder or schizophrenia, compared to key regulators of Alzheimer’s disease (BD vs. AD, 1 shared regulator, 1 expected by chance, p = 0.76; SCZ vs. AD, 13 shared regulators, 7 expected by chance, p = 0.02). Taken together, these results suggest that shared key regulator TFs connect diseases with overlapping clinical features.

We sought independent support for key regulator TFs by asking whether these TFs are associated with genetic risk for the same disease. We searched the GWAS catalog^56^ for genetic associations at loci containing TF genes, and we identified 13 instances in which a key regulator TF from our TRN model has been reported to be associated with genetic risk for the same disease (Fig. 2c; Table S8). This is twice as many instances of overlap between GWAS and TRN regulators as expected by chance (odds ratio = 2.05; p-value = 2.6e-2). TFs associated with both genetic and transcriptional changes in the same disease include key regulators for schizophrenia (*SREBF1*, *NPAS3*, *POU3F2*, *RFX2*, *KLF13*, *FOXN2*, *FOXO3*), bipolar disorder (*POU3F2*, *NPAS3*), and Alzheimer’s disease (*MEF2C*, *GLIS3*, *TFEB*, *NR3C2*). These results highlight the convergence of genetic risk and transcriptional changes on shared transcriptional regulatory networks in the brain.

### Disease-associated genetic variation influencing TF binding sites

The results above highlight putative *trans*-acting effects in which genetic changes at a small number of TF loci are linked to changes in the expression of many downstream target genes. We next sought to elucidate cis-acting network perturbations: genetic variants that alter a transcription factor binding site, perturbing a single edge in the network. eQTL analyses suggest that such effects are present at most genes, explaining hundreds of GWAS risk loci^4,9,57,58^.

To identify TFBS-disrupting SNPs, we intersected our model of human brain TFBSs with genetic variants from Kaviar^59^, focusing on SNPs that overlap TFBSs +/- 10 kb of the transcription start site for one of that TF’s predicted target genes. We identified 52,705 putative TFBS-disrupting SNPs (Supplementary Dataset 3). These SNPs are predicted to modify 67,152 of the 201,218 TF-target gene interactions in our TRN model (the number of modulated TF-target gene interactions is greater than the number of SNPs, since some SNPs overlap the binding sites for more than one TF and some TFs regulate more than one adjacent gene). These results support the idea that there are widespread effects of non-coding SNPs on gene regulation^60^.

To explore the potential impact of TFBS-disrupting variants on disease risk, we used our model to predict functional variants and target genes at risk loci for schizophrenia and Alzheimer’s disease. We found 17 schizophrenia-associated TFBS-disrupting SNPs (p < 1e-4), located at 11 of the 108 risk loci from the Psychiatric Genomic Consortium (Table S9). We also found four Alzheimer’s-associated TFBS-disrupting SNPs at four of the 19 risk loci from the International Genomics of Alzheimer's Project (IGAP; Table S10). 17 of these 21 disease-associated TFBS-disrupting SNPs are in strong linkage disequilibrium [LD] (r^2^ > 0.9) with an eQTL targeting the same gene. Notably, 11 of the 17 schizophrenia-associated TFBS-disrupting SNPs and two of the four AD-associated TFBS-disrupting SNPs are predicted to disrupt a binding site for one of the key regulator TFs for the same disease ‐‐ significantly more overlap than expected by chance (odds ratio = 3.3, p-value = 0.015). At SCZ risk loci, these included disrupted binding sites for the key regulators KLF15 (two binding sites), NR1H2, NR1H3, NR2F6, POU3F2 (two binding sites), PRRX1, RFX4, and SP3 (two binding sites) (Table 1; Table S9). At AD risk loci, these included disrupted binding sites for the key regulators KLF3 and KLF11. The enrichment for binding sites of key regulator TFs again highlights the convergence of genetics and gene expression on shared transcriptional networks.

**Table 1.** Binding sites for key regulator TFs are disrupted by SNPs associated with risk for schizophrenia. Schizophrenia-associated SNPs (p < 1e-4) that disrupted TF binding sites in the human brain were identified by scanning SNPs within 108 genome-wide significant schizophrenia risk loci^14^ for TFBSs from our TRN model. These loci contained 17 risk-associated, TFBS-disrupting SNPs, including 11 that involve key regulator TFs, shown in Table 1. For each SNP, the table shows the risk locus, as described by Ripke et al. (2014)^14^; the location of the risk-associated, TFBS-disrupting SNP; the p-value for the association of this SNP with schizophrenia, from Ripke et al. (2014)^14^; the key regulator TF whose binding site is predicted to be disrupted by this SNP; the predicted target gene, based on TRN analysis; and supporting evidence from eQTLs, compiled from HaploReg^64^ and the CommonMind Consortium^4^. Details of eQTL studies are shown in Table S9).

| SCZ Risk Locus | SNP | Postion | SCZ pval | TF | Target Gene | Distance to TSS | TF-gene Correlation (r) | eQTL pvalue |
| --- | --- | --- | --- | --- | --- | --- | --- | --- |
| chr2:57943593-58502192 | rs13384219 | chr2:58134458 | 2.1E-05 | *POU3F2* | *VRK2* | -336 | 0.67 | 2.0E-03 |
| chr2:198148577-198835577 | rs12618612 | chr2:198171058 | 3.7E-08 | *PRRX1* | *ANKRD44* | -4839 | 0.46 | 7.2E-18 |
| chr3:135807405-136615405 | rs9845788 | chr3:135914715 | 6.0E-07 | *SP3* | *MSL2* | -1371 | 0.49 |  |
| chr5:137598121-137948092 | rs154069 | chr5:137876920 | 2.3E-07 | *NR1H3*<br>*NR2F6* | *ETF1* | -2077 | 0.64<br>0.62 | 1.7E-10 |
| chr5:137598121-137948092 | rs11746692 | chr5:137946355 | 4.6E-07 | *NR1H2* | *CTNNA1* | -313 | 0.33 | 1.1E-04 |
| chr7:104598064-105063064 | rs62484724 | chr7:105049431 | 2.1E-05 | *POU3F2* | *SRPK2* | 9668 | 0.64 |  |
| chr10:104423800-105165583 | rs79780963 | chr10:104952499 | 7.5E-16 | *SP3* | *NT5C2* | -564 | 0.46 | 8.9E-57 |
| chr17:17722402-18030202 | rs7359509 | chr17:17741875 | 1.4E-06 | *KLF15* | *SREBF1* | 1546 | 0.67 | 9.4E-91 |
| chr17:17722402-18030202 | rs4925119 | chr17:17742904 | 5.9E-06 | *KLF15* | *SREBF1* | 2575 | 0.67 | 9.4E-91 |
| chr17:17722402-18030202 | rs6502618 | chr17:17746741 | 2.1E-06 | *RFX4* | *SREBF1* | 6411 | 0.50 | 9.4E-91 |

### Neuronal and microglia-enriched *MEF2C-regulated* networks associated with Alzheimer’s disease

Network modeling and genetic associations provided convergent evidence for a role of Myocyte-specific enhancer factor 2C (MEF2C) in Alzheimer’s disease. Specifically, our TRN model predicted that MEF2C regulates a module of neuronally-enriched target genes that are down-regulated in the DLPFC of individuals with AD (p = 1.1e-13). In addition, genetic variants at the *MEF2C* locus have been shown to be associated with risk for Alzheimer’s disease (p = 3e-8 in the IGAP study^61^).

To validate our network predictions for MEF2C, we studied publicly available RNA-seq from *Mef2c* conditional knockout mice. *MEF2C* has effects on both neuronal and microglial functions (e.g., neuroplasticity^62^ and inflammation^63^, respectively), and it has high levels of expression in both of these cell types^39^. We therefore considered expression changes in the brains of mice in which *Mef2c* was conditionally deleted either in neurons^62^ or in microglia (GSE98401).

Conditional deletion of *Mef2c* in neurons led to changes in the expression (p < 0.05) of 1,076 genes. Consistent with the predictions from our TRN model, these neuronal MEF2C target genes were enriched for genes that were down-regulated in AD cases vs. controls (p = 3.7e-4) and for neuron-specific genes (p = 4.4e-3). Also as predicted by our TRN model, neuronal targets of MEF2C were not enriched among genes that were up-regulated in AD cases vs. controls (p = 0.39). Neuronal MEF2C target genes were marginally over-represented for predicted MEF2C target genes in our TRN model (p = 0.05).

Conditional deletion of *Mef2c* in microglia led to changes in the expression (p < 0.05) of 896 genes. In contrast to neuronal MEF2C target genes, these microglial MEF2C target genes were not enriched among genes that were down-regulated in AD cases vs. controls (p = 0.96). Instead, microglial MEF2C target genes were enriched among genes that were up-regulated in AD cases vs. controls (p = 1.3e-6) and for microglia-specific genes (p = 9.4e-14). In addition, microglial MEF2C target genes did not significantly overlap the predictions from our TRN model (p = 0.28) or neuronal MEF2C target genes (p = 0.71).

Taken together, these results suggest that MEF2C contributes to both the down-regulation of neuronal transcripts and the up-regulation of microglial transcripts associated with Alzheimer’s disease. Neuronal effects of MEF2C were more accurately predicted by our TRN model, perhaps because microglia-specific gene networks are inactive in the healthy brain tissue used to train the TRN model^1^.

### An oligodendrocyte-enriched *SREBFl-regulated* network associated with schizophrenia

As with *MEF2C*, network modeling and genetic associations provided convergent evidence for a role of Sterol Regulatory Element Binding Transcription Factor 1 (*SREBF1*) in schizophrenia. Predicted target genes of SREBF1 in our TRN model were up-regulated in PFC from schizophrenia cases. To validate this finding, we used SREBF1 ChIP-seq data from the HepG2 and LoVo cell lines^36^. SREBF1 occupied a total of 3,438 genomic peaks in these two cell lines, located within 10kb of the transcription start sites for 3,909 genes. Although the ChIP-seq data are from non-neuronal cell lines, there was significant overlap between these experimentally characterized SREBF1 target genes and the predicted target genes from our TRN model (odds ratio = 1.5; p = 1.5e-3). In addition, SREBF1 target genes from ChIP-seq were enriched among genes up-regulated in PFC tissue of schizophrenia cases vs. controls across three independent cohorts^4,6,51^ (meta-analytic p-value = 8.0e-5). Therefore, SREBF1 ChIP-seq validate the prediction from our TRN model that SREBF1 is a key regulator of gene expression changes in schizophrenia.

The *SREBF1* gene sits at a genome-wide significant risk locus for schizophrenia at 17p11.2. The extent of LD at this locus spans eight genes: *ATPAF2*, *DRG2*, *GID4*, *LRRC48*, *MYO15A*, *RAI1*, *SREBF1*, and *TOM1L2*, so additional information is needed to identify the risk-associated gene (Fig. S3). Our model predicts three disease-associated, TFBS-disrupting SNPs at this locus ‐‐ rs7359509, rs4925119, and rs6502618 ‐‐ located 1, 4, and 6 kb upstream of *SREBF1*. Each of these SNPs disrupts a putative binding site for a TF that targets *SREBF1* in our TRN model, with eQTLs providing independent support for these SNP-gene associations (eQTL p-value = 9.4e-91^64^). Both rs7359509 and rs4925119 disrupt binding sites for the key regulator TF *KLF15*, while rs6502618 disrupts a binding site for the key regulator TF *RFX4*. Thus, eQTLs and TRN analysis suggest that *SREBF1* is most likely causal gene at this locus, and propose altered *KLF15* and *RFX4* regulation of *SREBF1* transcription as a mechanism.

*SREBF1* target genes were over-represented for oligodendrocyte-specific genes both in our TRN model (p-value = 3.2e-8) and based on ChIP-seq (p = 1.8e-3). Also, *SREBF1* target genes were enriched for the Gene Ontology terms “lipid binding” (p-value = 6.6e-5) and “phospholipid metabolism” (p-value = 2.1e-3). These enrichments are consistent with the well-established role of *SREBF1* in lipid biosynthesis^65^. We speculate that genetic perturbation of the (*RFX4-KLF15)-SREBF1-*(downstream target genes) network could contribute to deficits in lipid-rich myelin fibers (produced by oligodendrocytes), which have been described in the cortex of schizophrenia patients^66^.

### A neural stem cell‐ and astrocyte-enriched POU3F2-regulated network associated with bipolar disorder and schizophrenia

The transcription factor POU Class 3 Homeobox 2 (*POU3F2*, also known as *BRN2*) was implicated in bipolar disorder and schizophrenia via multiple network mechanisms: (i) *POU3F2* was a predicted key regulator of schizophrenia (p-value = 9.7e-6) and bipolar disorder (p-value = 5.2e-6) with predicted target genes over-represented among up-regulated genes in prefrontal cortex tissue from affected individuals with both diseases; (ii) The *POU3F2* genomic locus is associated with risk for bipolar disorder^15,17^; (iii) Predicted binding sites for *POU3F2* are disrupted by disease-associated SNPs at two genome-wide significant risk loci for schizophrenia (the *VRK2* and *SRPK2* loci). Thus, our TRN model implicates POU3F2 in schizophrenia and bipolar disorder both through c/s-acting effects of disease-associated variants that disrupt POU3F2 binding sites and through trans-acting effects on prefrontal cortex gene expression. In addition, *POU3F2* has well known roles in neural progenitors and neuronal differentiation^67^, suggesting a tractable, disease-relevant cell type in which to characterize its functions.

To validate a *cis*-acting effect of POU3F2, we characterized the regulatory impact of rs13384219 (hg19, chr2:58134458 A/G). This variant is located on one of several haplotypes at this locus that are associated with risk for both schizophrenia (Fig. 3a; lead SNP at this locus, p = 1e-11; rs13384219, p = 2.1e-5) and bipolar disorder^68^. rs13384219 is located 336 bp upstream of the transcription start site for *VRK2*. Our TRN model predicted a functional effect of this SNP on *POU3F2* binding for two reasons. First, the SNP overlaps a DNase I footprint spanning a sequence motif recognized by POU-domain transcription factors (Fig. 3b). Second, *POU3F2* expression was strongly correlated with *VRK2* expression in the Allen Human Brain Atlas (r = 0.67; Fig. 3c).

**Figure 3.**
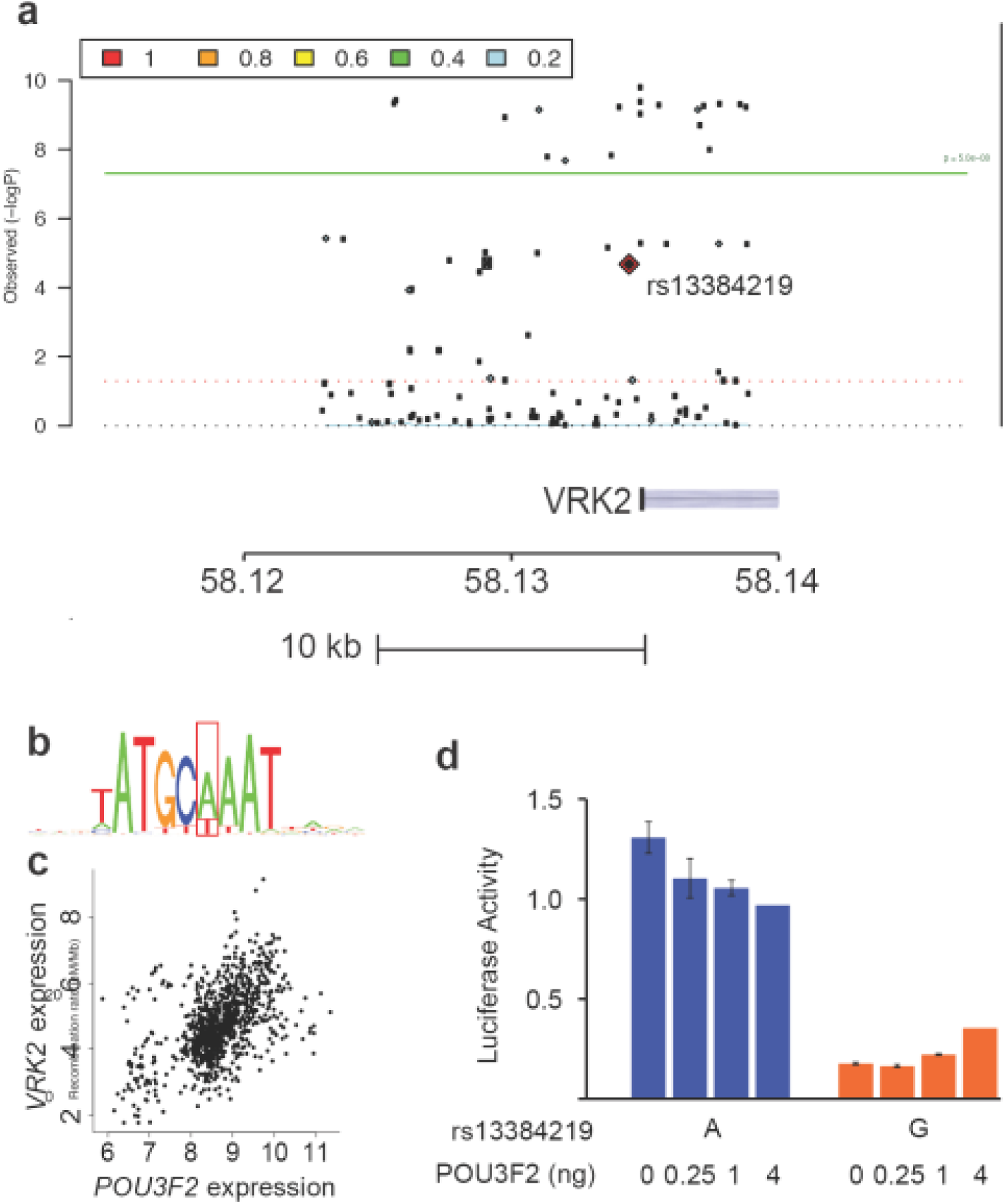
Modulation of a POU3F2 binding site by a schizophrenia-associated SNP in the *VRK2* promoter. **(a)** Region plot of the *VRK2* locus from GWAS of schizophrenia^14^. TRN analysis revealed a single risk-associated SNP at this locus (p < 1e-4) that overlaps a binding site for a TF that is a predicted regulator of VRK2, rs13384219. **(b)** rs13384219 disrupts a key residue in a sequence motif recognized by POU-domain TFs (the Pou2f2 motif is shown). **(c)** *POU3F2* expression is positively correlated with *VRK2* expression in the Allen Human Brain Atlas (r = 0.67) **(d)** Dual luciferase reporter assay comparing the activity of a *VRK2* promoter fragment containing either the A or G allele of rs13384219 in HEK293 cell. POU3F2 was overexpressed in combination with transfection of each luciferase construct to assess dose-dependent effects of POU3F2 and interactions with rs13384219 genotype.

We used luciferase assays to test the activity of a 436 bp fragment of the *VRK2* promoter containing either the A or G allele of rs13384219. The risk-associated G allele decreased the activity of the *VRK2* promoter in HEK293 cells (p = 4e-24; Fig. 3d). We repeated this experiment in combination with POU3F2 over-expression to test the effect of POU3F2 on the activity of the *VRK2* promoter fragment. *POU3F2* overexpression caused a dose-dependent decrease in the activity of the *VRK2* promoter fragment with the rs13384219 A allele. By contrast, POU3F2 over-expression caused a dose-dependent increase in the activity of the *VRK2* promoter fragment with the rs13384219 G allele (P_snp x pou3F2_ = 8e-4). These results validate our model’s predictions that POU3F2 regulates the *VRK2* promoter and that rs13384219 modifies this effect.

To validate trans-acting effects of POU3F2 and gain insight into its potential role in psychiatric disease, we overexpressed it in primary human neural stem cells (phNSCs; NSCs; Fig. 4). NSCs have low POU3F2 expression, and */n v/vo* induction of POU3F2 naturally occurs in early neural precursors, just subsequent to the stage of neurodevelopment at which our phNSCs were derived^67^. We collected RNA for microarray analysis at 3 and 10 days following infection and puromycin selection of cells transduced with a POU3F2 lentiviral expression vector. K-means clustering of these microarray data revealed nine gene clusters, two of which were differentially expressed upon POU3F2 overexpression specifically at the day 3 time point in all 4 replicate samples: cluster C6 showing repression and cluster C1 up-regulation (Fig. 5a).

**Figure 4.**
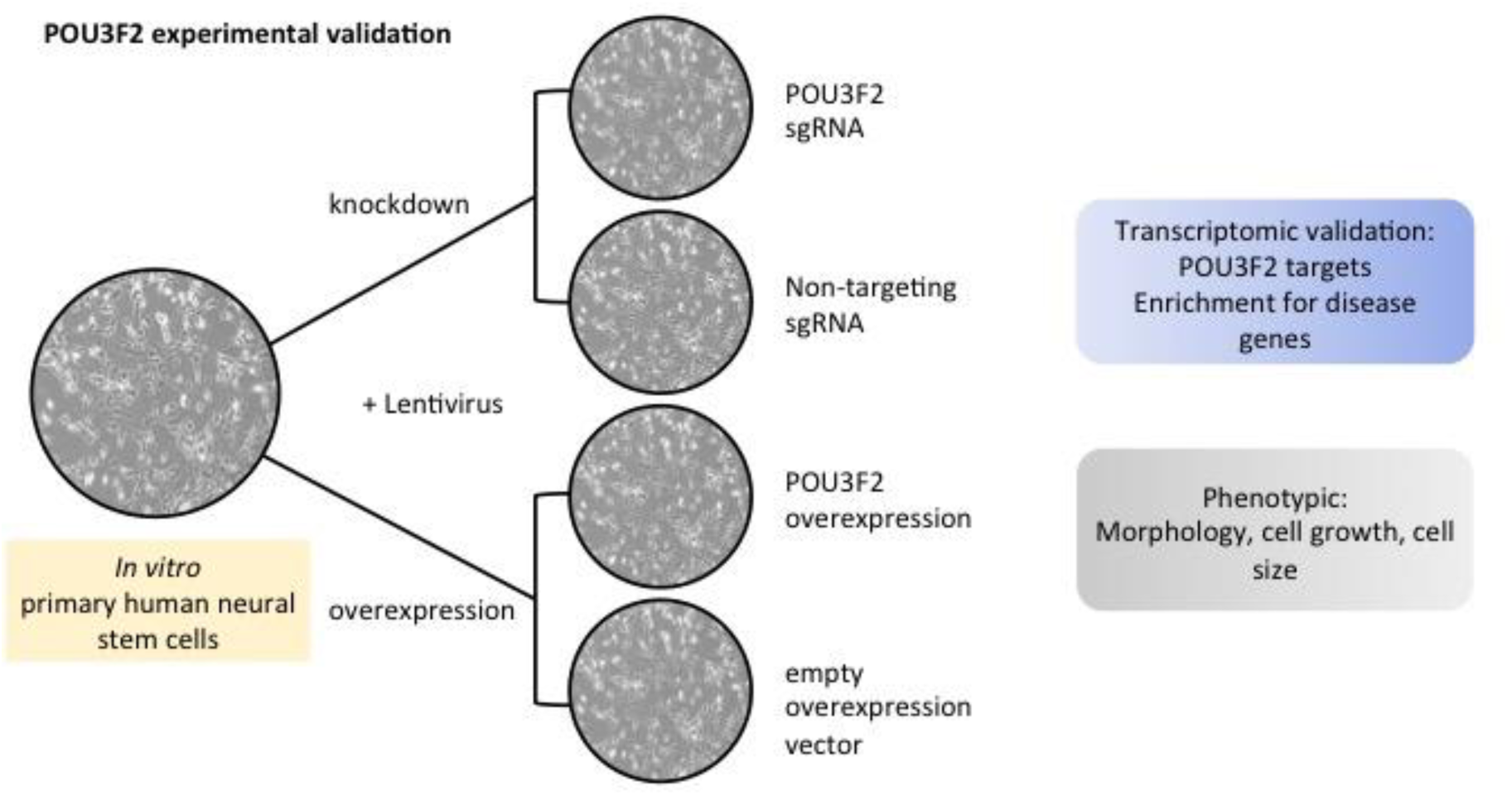
Schematic for experimental validation of POU3F2 gene targets in primary human neural stem cells.

**Figure 5.**
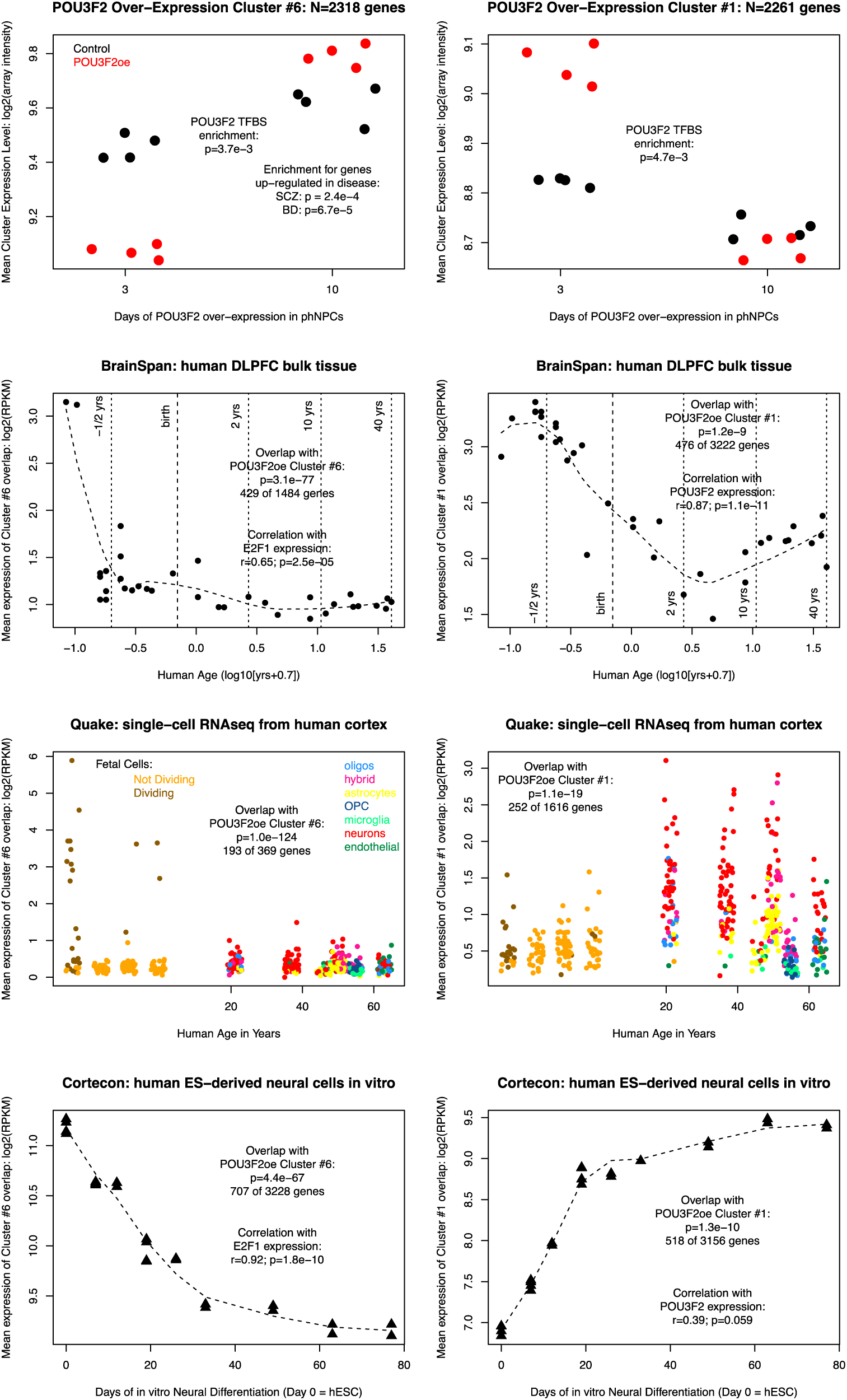
POU3F2 regulates a schizophrenia‐ and bipolar disorder-related gene network in neural stem cells. **We generated gene expression profiles** from primary human neural stem cells (phNSCs) after 3 and 10 days of transgenic overexpression of POU3F2 and from control phNSCs. We used K-means clustering to identify POU3F2-responsive gene clusters, then we compared these gene clusters to reference datasets from cortical development. **(a)** Mean cluster expression level (log2 array intensity) from genes in clusters C6 and C1 in POU3F2-overexpressing phNSCs and controls. **(b)** Mean expression (log2 RPKM) of genes in overlapping clusters from RNA-seq of developing dorsolateral prefrontal cortex from BrainSpan^69^. **(c)** Mean expression (log2 RPKM) of genes in overlapping clusters from single-cell RNA-seq of fetal and adult human cortex^71^; single-cell transcriptomes were assigned to brain cell types based on cell type markers and clustering, as described by the study’s authors. **(d)** Mean expression (log2 RPKM) of genes in overlapping clusters from *in vitro* differentiation of human ES cells into forebrain neurons.^92^

We compared genes in clusters C6 and C1 to our TRN model to evaluate the overlap with our *in silico* network predictions. Genes in C6 were enriched for genes whose promoters and proximal enhancers (+/-10 kb) harbor multiple brain-occupied POU3F2 binding sites from our model (p = 3.7e-3), as well as for computationally-predicted *POU3F2* target genes in the adult brain (p = 8.5e-4, odds ratio = 1.44, 105 shared genes). Importantly, genes in C6 were over-represented among genes that were up-regulated in prefrontal cortex from individuals with schizophrenia (p = 2.4e-4) and bipolar disorder (p = 6.7e-5). Genes in C1 were also enriched for genes whose promoters and proximal enhancers harbor multiple brain-occupied POU3F2 binding sites from our model (p = 4.7e-3). However, the genes in C1 did not overlap computationally-predicted POU3F2 target genes or genes up-regulated in schizophrenia or bipolar disorder, suggesting that these POU3F2 target genes are either non-targeted in the adult brain or are missed by our computational model. Therefore, POU3F2 overexpression in phNSCs validated our network prediction that POU3F2 is a key regulator of gene expression changes in prefrontal cortex from individuals with schizophrenia and bipolar disorder and identified additional target genes that may be specific to neurodevelopment.

To gain perspective on how these expression changes relate to *in vivo* development, we explored the expression patterns of genes from clusters C6 and C1 in several other neurodevelopmental gene expression datasets: (i) RNA-seq of dorsolateral prefrontal cortex development from BrainSpan^69^; (ii) gene expression microarrays of laser-captured regions of the developing macaque cortex^70^; (iii) single-cell RNA-seq of the developing human cortex^71^; and (iv) RNA-seq of *in vitro* differentiation of human induced pluripotent stem cells toward neurons^72^.

Genes in C6, down-regulated upon POU3F2 overexpression, were enriched for genes involved in the cell cycle (Gene Ontology, p=5.8e-75). Genes in this cluster overlapped significantly with clusters obtained from *in vivo* neocortical data, showing high expression during the first trimester of *in utero* development and low expression thereafter, in both human and non-human primate (Fig. 5b, Fig. S4). TRN analysis of the genes in both cluster C6 and the overlapping human DFC cluster indicated the involvement of several TFs known to control cell cycle, including the top hit E2F1 (p=1.3e-32), which shows high correlation with the DFC cluster itself (r=0.65, p=2.5e-5). Single-cell data from the developing human neocortex indicate that these genes are expressed specifically in dividing cells of the prenatal neocortex (Fig. 5c). Remarkably, a subset of the genes in the C6 cluster, including GLI3, are also highly expressed in astrocytes of the adult cortex (Fig. S5).

Genes in C1, up-regulated upon POU3F2 overexpression, were enriched for genes involved in transcription (Gene Ontology, p=4.1e-13). These genes overlapped a cluster of genes whose neocortical expression peaks in the middle of the second trimester and which correlates with POU3F2 expression itself (Fig. 5b, r=0.87). Single-cell data suggest that these genes are expressed in cells of the developing neocortex that are not dividing and that these putative targets of POU3F2 transcriptional activation increase further as neurons mature (Fig. 5c).

These observations suggest that POU3F2 overexpression in primary human phNSCs *in vitro* recapitulates an *in vitro* cellular transition out of an early highly proliferative state and into a subsequent POU3F2-driven state during the second trimester of human neocortical development. Similar cluster overlapping with data from the CORTECON project^60^ indicates that the dynamics of these gene clusters are robust in distinct *in vitro* neural differentiation protocols (Fig. 5d).

To directly test the effects of POU3F2 on the transition from proliferative to non-proliferative cell states, we quantified cell proliferation, cell size, and nucleus size in phNSCs and in cells differentiated for two weeks toward neurons or astrocytes. We overexpressed POU3F2 as in the gene expression profiling experiments above, and we knocked it out via lentiviral delivery of a CRISPR/Cas9 sgRNA. We observed a significant decrease in proliferation in POU3F2 overexpressing phNSCs compared to control phNSCs (̃2 fold decrease, p=0.0043, Fig. 6a). There was a trend toward increased proliferation in POU3F2 knockout phNSCs, but this was not significant (p=0.1523). In addition, we observed a significant increase in cell size in POU3F2-overexpressing cells compared to controls, both in phNSCs (13 fold increase, p<0.0001) and in cells differentiated toward cortical neurons for two weeks (̃6 fold increase, p<0.001) (Fig. 6b,c)). We also observed a significant increase in nucleus size in POU3F2 overexpressing cells in the NSC state (̃1.5 fold increase, p=0.01). These findings suggest that POU3F2 overexpression inhibits cellular proliferation, possibly via its repression of cell cycle genes and of genes expressed in proliferative phNSCs.

**Figure 6.**
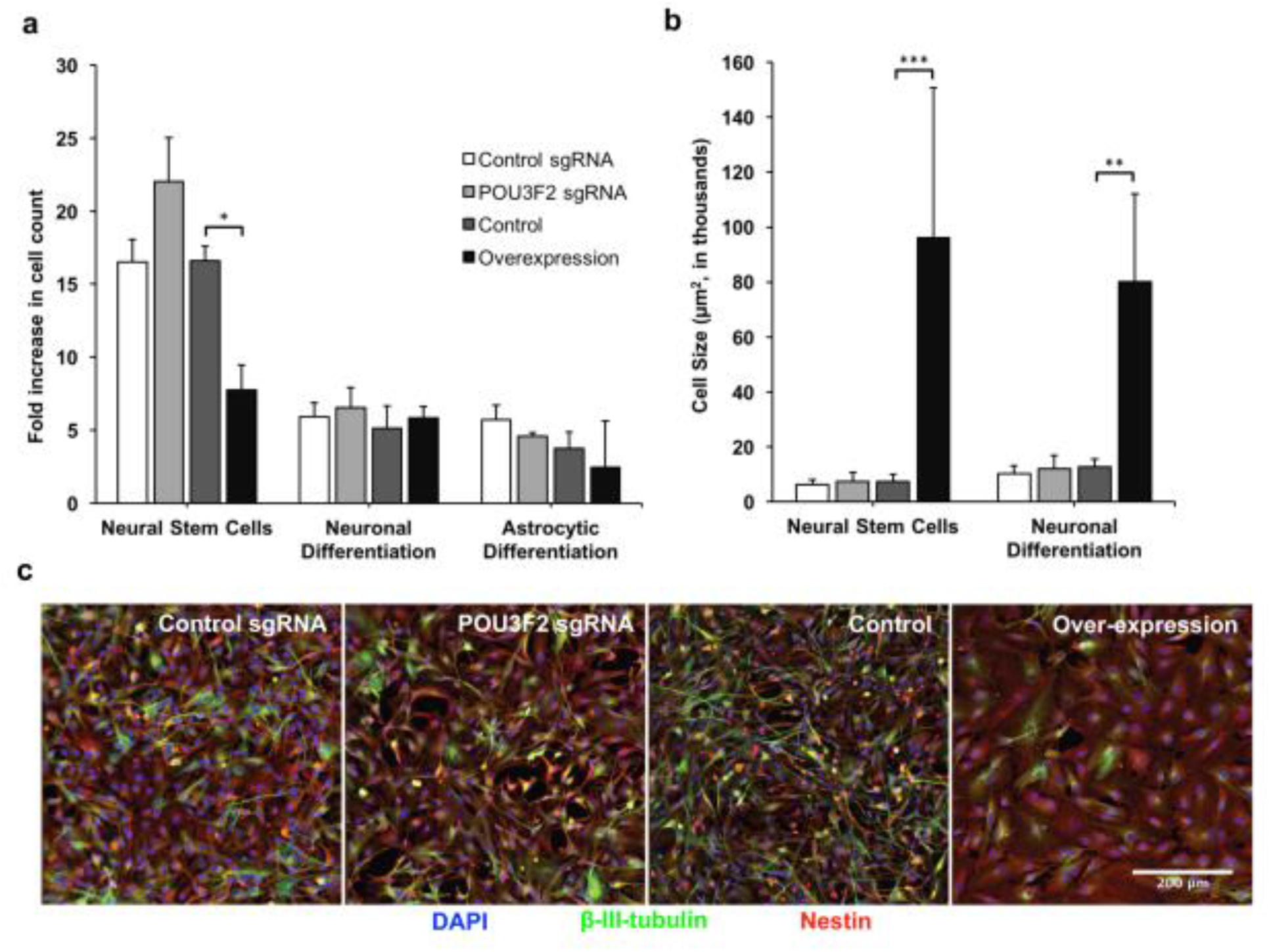
POU3F2 represses cellular proliferation in primary human neural stem cells. We perturbed POU3F2 expression in primary human neural stem cells (phNSCs) by knocking it out with a CRISPR/Cas9 sgRNA or with transgenic over-expression, then assessed effects on cellular prolifersation. (a) POU3F2 overexpression decreased cellular proliferation in phNSCs to levels comparable to proliferation in cells differentiated toward neurons or astrocytes. POU3F2 sgRNA did not significantly influence proliferation. (b) POU3F2 overexpression increases cell size, both in phNSCs and in cells differentiated toward neurons. (c) Immunofluorescence images of phNSCs and of differentiated neurons, showing effects of POU3F2 overexpression on cell size and morphology.

## Discussion

In this study, we have presented a genome-scale transcriptional regulatory network model for the human brain. We identified key regulator TFs that are implicated in gene regulatory changes underlying psychiatric and neurodegenerative disorders, involving changes in gene expression as well as putative *cis-* and trans-acting genetic variation. We systematically validated these mechanisms through comparison to ChIP-seq and single-gene perturbation experiments. We describe convergent genetic and transcriptional network support for roles of key regulator TFs in multiple diseases, including the MEF2C-regulated network in Alzheimer’s disease; the SREBF1-regulated network in schizophrenia; and the POU3F2-regulated network in bipolar disorder.

Genetic associations and transcriptional changes in a disease often converged on the same key regulator TFs. Convergence on shared networks provides independent support for the associations of these TFs with disease. In addition, these results support the view that transcriptional network changes are a core feature underlying many human diseases. Recently, Boyle et al. (2017)^73^ invoked the small-world property of biological networks - nearly all genes in a cell form a single interconnected network of direct and indirect connections ‐‐ to explain how virtually any gene expressed in a disease-relevant cell type could contribute to genetic risk for a disease. Our results build on this insight, yet emphasize that an understanding of network structure can reveal a relatively small number of core genes.

Our models identify key regulators of psychiatric and neurodegenerative diseases acting in multiple brain cell types. In Alzheimer’s disease, neuronally-enriched TF networks were often down-regulated, whereas microglia-enriched TF networks were up-regulated. Remarkably, MEF2C may contribute to both of these processes. Most research on schizophrenia and bipolar disorder has focused on neuronal mechanisms. Our results suggest that neuronal gene expression changes in these diseases are mediated by TFs such as *MEF2A*, a key regulator TF in our model whose target genes were enriched in neurons (Table S3) and which has previously been shown to influence synaptic function^74^. Strikingly, many disease-perturbed networks in psychiatric disorders were enriched in non-neuronal brain cell types. For instance, the oligodendrocyte-enriched *SREBF1* network was implicated by both genetic and gene expression changes.

Strikingly, several key regulator TFs for schizophrenia and bipolar disorder are well known for their roles in neural stem cells, including *HES1*, *MEIS2*, *NKX2-1*, *NPAS3*, RFX2, *RFX4*, *SOX2*, and *SOX9*, among others^38^. Of these, *POU3F2*^15,17^, *SOX2*^14,21^, NPAS3^18^,^75^, and *RFX4*^*76*^ are also associated with genetic risk for schizophrenia or bipolar disorder, suggesting an etiological role. Interestingly, we identified these NSC-related key regulator TFs based on differential expression in the adult brain. Most of these TFs are also highly expressed in adult astrocytes, and their target genes in our TRN model were enriched for astrocyte-specific genes (Table S3). These pleiotropic effects make it difficult to discern the time points and cell types in which these TFs influence psychiatric disorders. A neurodevelopmental hypothesis is compelling, since adverse events during fetal development are among the strongest non-genetic risk factors for schizophrenia and mood disorders and many schizophrenia risk genes are most highly enriched during fetal brain development^77,78^. Changes in adult astrocytes likely also contribute to psychiatric disorders. Several of these NSC/astrocyte TFs have been implicated in cancer stem cells within astrocytic tumors^79^ and in de-differentiation of astrocytes and adult neurogenesis^80^, suggesting functions in proliferative astrocyte states. Understanding how disease risk emerges from changes that occur across all of these cell types and time points remains an exciting future challenge.

Convergent lines of evidence led us to focus on POU3F2 as a central regulator of gene expression changes in schizophrenia and bipolar disorder. We validated key network predictions that POU3F2 target genes are over-represented among differentially expressed genes in prefrontal cortex from schizophrenia and bipolar disorder vs. controls and that a risk-associated SNP near *VRK2* influences gene expression through an interaction with POU3F2. We show that POU3F2 represses cell cycle genes and regulates the proliferation of neural stem cells. These anti-proliferative effects are consistent with two recent reports linking *POU3F2* to neural proliferation phenotypes in stem cell models of autism^81,82^. This manuscript is being submitted in parallel with a second paper demonstrating that POU3F2 regulates SCZ-related gene expression changes in prefrontal cortex. The two studies converged on this conclusion independently, despite utilizing different transcriptomic datasets and network reconstruction approaches. We exchanged gene lists and found that there is significant overlap between the lists of computationally predicted POU3F2 target genes described in the two studies (p-value = 4.5e-7, odds ratio=2.4). Therefore, our shared conclusions represent a true convergence of our approaches on the same disease-perturbed network. This convergence across studies emphasizes the reproducibility of results in systems biology and the importance of POU3F2 as a key regulator of bipolar disorder. Identifying genes involved in bipolar disorder has been particularly difficult: 30 genome-wide significant risk loci reported in the largest GWAS to date^68^, but biological follow-up experiments have been undertaken for only a few positional candidate genes at these loci^83,84^. The convergent evidence presented here adds POU3F2 to a very short list of well-supported bipolar disorder risk genes.

Our TRN model is a broadly applicable resource for future genetic and genomic studies of the human brain and of brain diseases. Our models of binding sites and target genes for 741 TFs, as well as predictions of TFBS-disrupting SNPs, are available online (http://amentlab.igs.umaryland.edu/psych-trn-pfc2016/). These resources can now be used to identify key regulators of gene expression changes in any brain-related transcriptomics experiment and for functional analysis of non-coding genetic variation. As such, this study presents a roadmap for understanding brain gene regulation in human health and disease. The same approach can also be taken for diseases impacting other organs, using a wide range of publicly available epigenomic and transcriptomic data.

## IV. METHODS

### DNase-seq of human brain tissue

We used publicly available data from 15 DNase-seq experiments with human brain tissue, generated by the ENCODE project (Table S1). FASTQ files were downloaded from https://www.encodeproject.org in January, 2016. Note that ENCODE makes a distinction between sample and experiment in that one experiment can contain more than one sample. An experiment can contain both single or paired-end reads, with varying depth of sequencing and varying read length in a single experiment. https://www.encodeproject.org/search/?type=Experiment&assay_slims=DNA+accessibility&assay_title=DNase-seq&award.project=ENCODE&replicates.library.biosample.donor.organism.scientific_name=Homo+sapiens&organ_slims=brain

### Alignment

For each DNase-seq experiment, we started with the FASTQ files available on https://www.encodeproject.org/. Each FASTQ file (or paired-end files) was aligned to GRCh38 using the SNAP algorithm^24^. SNAP creates a large hash table of indices prior to alignment with a default seed length of 20. Because most of the FASTQ files were short ( < 50 bases), we reduced the seed size to a length of 16. The number of footprints found from an experiment is strongly correlated to the depth of sequencing. As the length of reads was variable, the number of mismatches allowed was adjusted accordingly as a parameter of SNAP.

### Identifying regions of open chromatin

To identify regions of open chromatin from the aligned BAM files, we used F-seq^25^. Our choice of F-seq was based on work from Koohy *et al*.^85^, who compared four different approaches (F-seq, Hotspot, MACS and ZINBA) on DHS data. We utilized the parameters they found to produce the best results in their benchmark comparison to ENCODE ChIP-seq data, using a minimum length of 400 bases.

### Running Wellington

Using the output bed files from F-seq, we identified putative footprints using Wellington^26^. Wellington was run with standard settings and ‐fdrlimit set to ‐1. Instructions for running Wellington can be found here: http://pythonhosted.org/pyDNase/tutorial.html.

### FIMO Database

All potential motif matches in GRCh38 were identified using FIMO^86^ and parsed using customized bash scripts. As an input to FIMO, we used JASPAR CORE Vertebrate 2016 collection (http://jaspar.genereg.net/html/DOWNLOAD/). and additional motifs from the HOMOCOMO and Swiss Regulon databases that match TF families not included in JASPAR CORE. Motif point-weight matrices were downloaded with the MEME suite^87^. The footprints identified from Wellington were intersected with the FIMO catalog. To maximize coverage, and because of the potential imprecise nature of footprints, if any part of a known motif overlapped with a single base of the footprint, an entry was created. We assigned motifs to their cognate TFs, as well as to additional TFs in the same DNA-binding domain family from the TFClass database^88^.

### Transcriptional regulatory network model for the human brain

We predicted TF-target gene interactions by integrating brain-specific DNase-seq genomic footprints with human brain gene expression profiles from the Allen Brain Atlas (http://human.brain-map.org/static/download). The Allen Brain Atlas consists of 2,748 microarray gene expression profiles of microdissected tissue from six human brains. We applied two gene expression-based methods to predict active TF-target gene interactions among these candidate regulators: (1) Pearson correlation and (2) LASSO regression (see below for details). We constructed separate models using the gene expression data from each of five brains in the ABA, leaving the data from the last of the six Allen Brain Atlas brains as a test set. Finally, we created a consensus model, retaining TF-target gene interactions that were selected by both the Pearson and LASSO methods in at least two of the five brains.

#### Selection of candidate regulators based on genomic footprinting

We counted the number of footprints for each TF in proximal and distal regions around each gene’s canonical transcription start site (TSS; ENSEMBL gene models), with proximal regions defined as +/- 10kb from the TSS and distal regions defined as +/- 1Mb from the target gene’s TSS and >10kb from any gene’s TSS. TFs with low-complexity sequence motifs generally have large numbers of motifs. To reduce the bias of our models toward TFs with low-complexity motifs, we quantile normalized the footprint counts for each TF across all genes. We selected as candidate regulators those TFs that had normalized footprint counts in the upper quartile, in either the proximal or distal region around each gene’s TSS.

#### Methods to predict active TF-target gene interactions

1. <u>Pearson correlation</u>. We calculated the Pearson correlation (r) between each TF-gene pair, across the samples from each of the six ABA brains, separately. TF-gene pairs with |r| > 0.25 were considered correlated.
2. <u>LASSO regression</u>. We fit a LASSO regression model to predict the expression of each target gene based on a linear combination of the expression patterns of its candidate regulators. These LASSO regression models were fit using the glmnet R package^89^. We considered candidate regulators with proximal binding sites and with distal binding sites, sequentially. First, we constructed a model considering only TFs whose binding sites were enriched in proximal regions (+/-10kb from the TSS). We then used the predictions from this model as an offset in a second model in which we considered TFs whose binding sites were enriched only in distal regions (>10kb and <1Mb from the TSS).

### Over-representation of TF target genes among differentially expressed genes, cell-type specific genes, functional categories, and curated gene sets

We tested for over-representation using one-tailed hypergeometric distributions. We calculated a False Discovery Rate for each test using the Benjamini-Hochberg method.

### Prediction of disease-associated TFBS-disrupting variants

Using bedtools, we intersected TFBSs with genetic variants from the Kaviar database (http://db.systemsbiology.net/kaviar/), restricting our analysis to variants in dbSNP. We included all variants that overlapped a predicted TFBS by at least 1 bp. Next we intersected this table of putative TFBS-disrupting variants with summary statistics from the Psychiatric Genomics Consortium GWAS of schizophrenia^14^ (https://www.med.unc.edu/pgc/results-and-downloads) and from the International Genomics of Alzheimer’s Project (http://web.pasteur-lille.fr/en/recherche/u744/igap/igap_download.php). For schizophrenia, we studied 108 predefined risk loci^14^. For Alzheimer’s disease, we calculated the bounds of risk loci from GWAS summary statistics using the ‐‐clump and ‐‐ld-snp-list commands in PLINK, as described^14^. We selected TFBS-disrupting variants that were associated with schizophrenia risk at p-value < 1e-4 and located within the extent of LD at each genome-wide significant locus.

### Neural Stem Cells

NSC lines were grown in neural expansion media (NEM) supplemented with EGF and FGF-2 (20 ng/ml) (PeproTech) on laminin (Sigma L2020) ‐coated polystyrene plates and passaged as previously described^90^. Neural expansion media (NEM): NeuroCult NS-A Basal Medium (Stem Cell Technologies), DMEM/F12, Antibiotic-Antimycotic, GlutaMax, B-27 supplement, N-2 supplement, 1:1000 EGF, and 1:1000 FGF.

### Lentivirus Infections

VSV-G pseudotyped, self-inactivating lentivirus was prepared by transfecting 293T cells with 1.5 μg pVSV-G, 3 μg psPAX-2, and 6 μg pRRL lentiviral vectors - either lentiCRISPR or lenti-SFFV-POU3F2-Myc-DDK. Lentiviral supernatants were collected 48 hours later and transferred to human neural stem cells dishes (MOI = ̃ 1). Positively transduced cells were selected with 0.6 ug/mL puromycin for 3 days. Gene editing was evaluated using the Surveyor^®^ Assay (Integrated DNA Technologies). Overexpression was evaluated using qRT-PCR and Western blotting.

### Generation and Analysis of CRISPR-Cas9 mutants

*POU3F2* was edited in neural stem cells using a custom guide site targeting the following sequence: ATCGTGCACGCCGAGCCGCCCGG. Genomic DNA was extracted from transduced and puromycin-selected cells (*POU3F2* targeted and non-targeting guide site control) using Thermo genomic DNA purification kit (K0512) and amplified using AccuPrime *Taq* DNA polymerase. The following primers were used to amplify a 431 bp region framing the target site for analysis of editing using the Surveyor^®^ Mutation Detection Kit (IDT #706025).

POU3F2_For: AGAGCGAGAAGGAGGGAGAG

POU3F2_Rev: GTGATCCACTGGTGAGCGTG

Four microliters of PCR product was used for TOPO cloning for sequencing (Thermo #K457501) and transformed into TOP10 cells. 34 colonies were picked, grown up, and plasmid DNA extracted (Qiagen Miniprep Kit). Inserts were sequenced at GeneWiz, and percentage gene editing in cell population was determined from alignment to reference sequence. Results from these validation experiments are shown in Fig. S6 and Table S10, indicating ̃50% editing efficiency.

### Western Blot Analysis

We used Western blots and qPCR to ePCR to evaluate POU3F2 overexpression. Cells were harvested following lentiviral infection and selection, washed with PBS, and protein extracted using RIPA buffer on ice for 20 minutes. 30 ul of Nupage LDS buffer was added to 100 ul of sample (in RIPA) and boiled at 80 degrees C for 10 min. 12 microliters was loaded onto 4-12% Bis-Tris gel and run for 1 hour at 180 volts. iBlot transfer system was used to transfer to PVDF membrane, according to manufacturer’s instructions. Membrane was cut between 51 and 39 kDa bands indicated by See Blue Plus 2 pre-stained ladder. The following commercial antibodies were used: POU3F2 (Abcam ab137469, 1:1000), FLAG (Sigma #F1804, 1:5000), and GAPDH (Abcam ab37168, 1:5000). A FluorChem was used to image the blots using SuperSignal West Dura chemiluminescent substrate (Thermo #34075). Results are shown in Fig. S7.

### qPCR

Total RNA was extracted using Qiagen miRNeasy kit. cDNA was reverse transcribed using the VILO mastermix kit (Thermo #11755050). A multiplex qPCR assay was run in triplicate for each sample, from duplicate RNA extractions of each modified or control NSC cell line. Multiplex reactions contained SsoFast Universal Probes mix, POU3F2 assay (FAM; IDT custom primer/probe assay) and ACTB assay (HEX; IDT #Hs.PT.56a.40703009). Samples were run on BioRad CFX96 Real-Time system and quantified using the CFX software. Results are shown in Fig. S8.

### Neural stem cell differentiation

Protocols for NSC differentiation toward astrocytes and neurons were modified from previously reported methods^91^. Following cell count, 15,000 cells were seeded into each well on 4-well chamber slides (Thermo #154526PK). Cells were treated with either neuronal-differentiation media (NeuroCult NS-A Basal Medium Stem Cell Technologies, 2% B-27 Serum-Free Supplement, 1% GlutaMax, 1% Antibiotic-Antimycotic) or astrocyte differentiation media (DMEM L-glutamine, high glucose, 1% N-2 supplement, 1% GlutaMax, 1% FBS). Control cells were maintained in NEM. Media was changed every 3-4 days over an 18-day differentiation period. After one week, 0.05 mM dibutyryl cAMP was added to neuronal differentiation media for the remaining differentiation.

### Immunofluorescence assays

Cells were fixed for 10 minutes in warm 4% paraformaldehyde. Immediately following fixation, standard IF protocol (Cell Signaling Technologies) was conducted with citrate buffer antigen retrieval. Both primary and secondary antibodies were allowed to incubate at 4 degrees overnight. To detect NSC, neuronal, and glial markers we used Nestin (R&D anti-hNestin purified mouse monoclonal IgG #5568P), b-III-tubulin (CST beta3-Tubulin Rabbit mAb #5568P), and GFAP antibodies (CST GFAP (GA5) mouse mAb #3670S), respectively. Secondary antibodies were purchased from Cell Signaling Technologies (Anti-Mouse IgG - Alexa Fluor 647 conjugate #4410, Anti-Rabbit IgG - Alexa Fluor 488 conjugate #4412, and Anti-Rabbit IgG - Alexa Fluor 555 conjugate #4413).

### Quantification of IF images for cell proliferation and size analysis

Cells were stained for DAPI (CST 4063S), stem cell marker Nestin (R&D MAB1259), and neuronal marker b-III-tubulin (CST 5568P). Image J software (FIJI) was used for all analyses. Images for quantifying cell size were taken on a Leica DM IRBE with a 40X NA 1.25 Oil Immersion objective at University of Washington’s W.M. Keck Microscopy Center. A total area of 0.553 mm^2^ per cell line per condition was visualized by combining 18 individual images within the imaging program MetaMorph. Cell size was quantified by tracing and recording the area of a randomly selected subset of cells after immunofluorescence staining using the marker that best captured the entire cell (Nestin for NSC state, b-III-tubulin for neuronal differentiation). Cell size was not quantified for astrocytic differentiation due to poor resolution of cell boundaries. Cell proliferation was quantified by using DAPI to count the number of nuclei within an image of each well that was 2.5% of the total well (4.6 mm^2^ sampled of 1.7 cm^2^ total well area). Images for quantifying cell size were taken on a Leica DM IRBE with a 10X objective at University of Washington’s W.M. Keck Microscopy Center. We calculated the fold change in number of cells by extrapolating the number of nuclei in the imaged area to the number of cells total in each well. Finally, given the seed density, the fold increase was quantified.

### Microarrays

Total RNA was extracted using the miRNeasy extraction kit (Qiagen) at three days and ten days post-puromycin selection of lentiviral transduced hNSCs. Two individual RNA extractions were performed from each plate at each time point, and each viral construct was transduced into two plates of cells. Gene expression was quantified using SurePrint G3 Human Gene Expression 8x60K v2 Microarray (#G4851B). Samples overexpressing POU3F2 were compared to samples that were transduced with a control vector that did not contain the *POU3F2* coding sequence. Microarray data have been deposited in the Gene Expression Omnibus (GSE102122; reviewer token klifsmswxfsftob).

### Gene Expression Analysis

In Figure 5, we used the intersectoR() function in the projectoR package in the R statistical language (https://github.com/genesofeve/projectoR) to explore gene membership overlaps between the gene clusters defined in the POU3F2 over-expression experiments we performed here and other key public data sets relevant to brain development and neuropsychiatric disease. This uses a hypergeometric test, phyper(), to determine the statistical significance of the number of genes shared across clusters in the different experiments. Correlation between individual genes’ expression and cluster averages were determined using the cor.test() function. Enrichment of disease gene DEG lists and genes annotated to particular GO terms was calculated using the geneSetTest() function in the limma package within Bioconductor.

### Luciferase Reporter Assay

*VRK2* promoter sequences were amplified from the genomic DNA of an individual whose genome was heterozygous at this position based on whole-genome sequencing. Amplified DNA was cloned into the pGL4.10[*luc2*] reporter plasmid (Promega #E6651) upstream of the *luc* transcription start site. Reporter constructs were co-transfected with a pRL-CMV Renilla vector (ratio of 100 ng luciferase: 5 ng renilla control plasmid) into HEK293 cells using Lipofectamine 2000. After 48 hrs, cells were harvested and luciferase and renilla activity was assayed on a Synergy H4 Plate Reader using the Dual-Luciferase^®^ Reporter Assay (Promega #E1910). All reported values were normalized to renilla co-transfection controls. All experiments were performed with 3 biological replicates per condition in a 24-well plate format. Results are representative of at least 2 independent experiments. Barplots are presented as mean +/- s.d.

## AUTHOR CONTRIBUTIONS

SAA and JRP designed the experiments. JRP, BB and DB performed the experiments. SAA, JRP, DB, CF, RO, and CC analyzed the data. SAA, JP, and CC wrote the paper. All authors contributed to editing the manuscript.

## ACKNOWLEDGMENTS

Patrick Paddison, Yu Ding and Chad Toledo (Fred Hutchinson Cancer Research Cneter) provided human neural stem cells. Elizabeth Gray, Daniel Stetson, and Kathleen Pestal (University of Washington) provided LentiCRISPR plasmids and protocols. John Kelsoe and Tatyana Shekhtman (University of California, San Diego) provided genomic DNA for promoter cloning of *VRK2*. Nathaniel Peters (W.M. Keck Microscopy Center, University of Washington) provided immunofluorescence imaging and advice. Victor Felix maintains resources available at amentlab.igs.umaryland.edu. Gene Robinson provided helpful discussion and comments on an early version of this manuscript. This work was supported by a National Science Foundation Graduate Research Fellowship (JRP); a NARSAD Young Investigator Award from the Brain and Behavior Research Foundation (SAA), the NIGMS Center for Systems Biology at the Institute for Systems Biology (P50 GM076547; LH, NDP), and by the Big Data for Discovery Science Center of the NIH Big Data to Knowledge program (NDP, LH).

---

**Figure S1.**
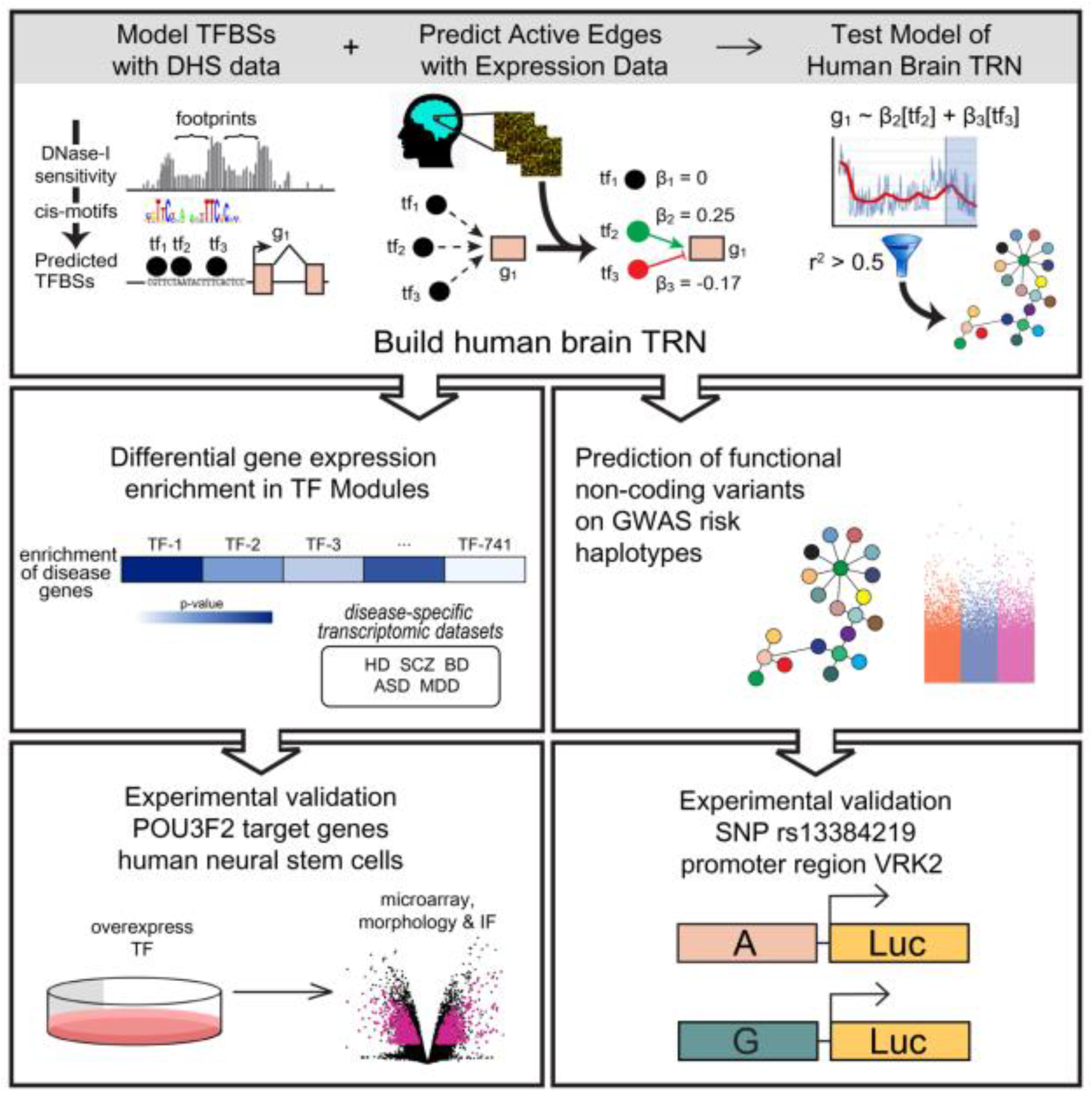
Graphical abstract.

**Figure S2.**
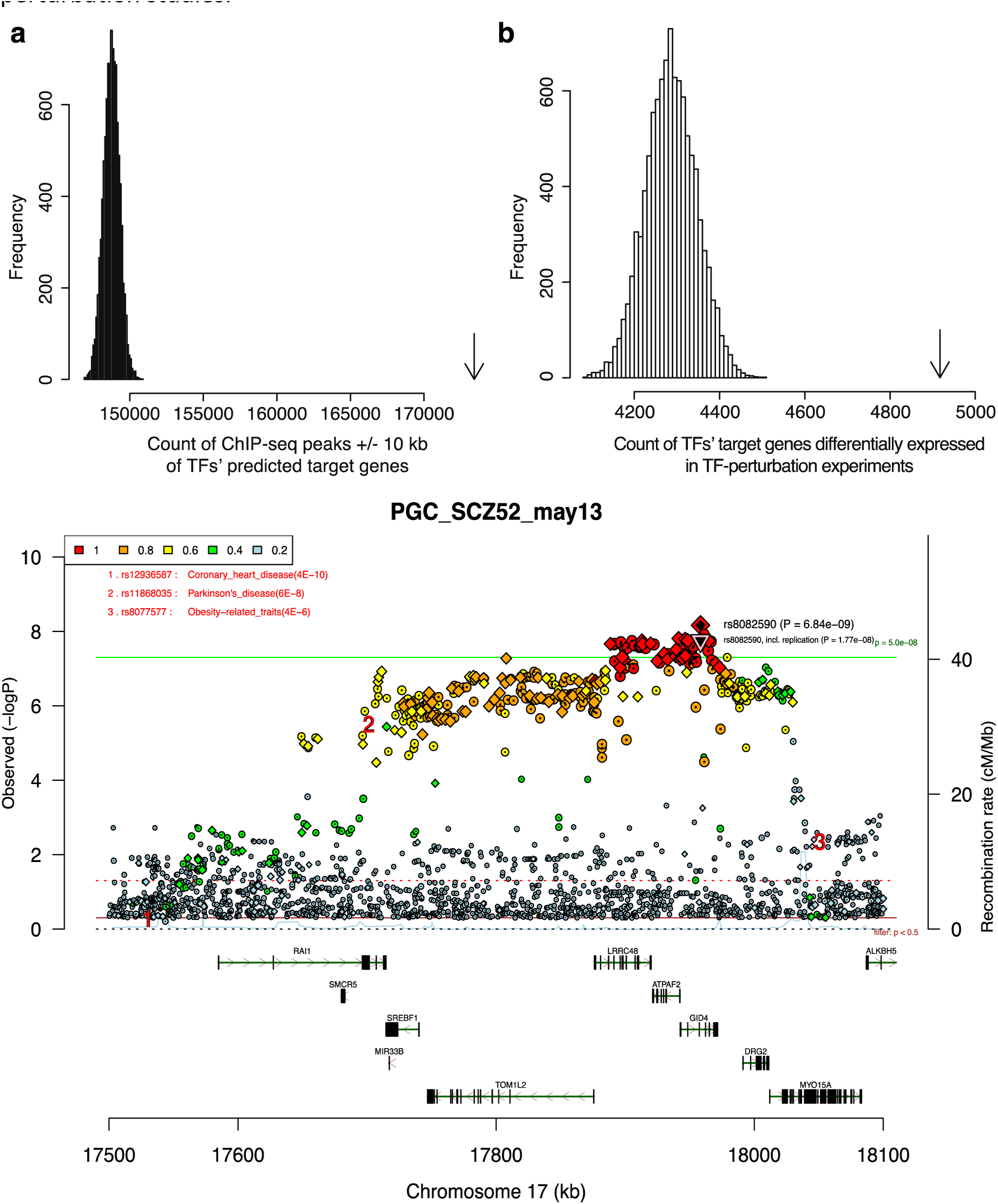
Computationally predicted TF-target gene interactions are enriched for experimentally characterized TF-target gene’s from published ChIP-seq and single-gene perturbation studies. A. Comparison of computationally predited TF-target gene interactions to ChIP-seq of 402 human TFs from the GTRD database. X-axis indicates the total number of instances in which a ChIP-seq peak from GTRD in located within 10kb of the TSS for a predicted target gene of that same TF. Arrow indicates the observed count from the TRN model. Histogram indicates an empirical null distribution derived from 10,000 permutations of edges from the brain TRN model. The observed value was never achieved in 10,000 permutations. B. Comparison of computationally predited TF-target gene interactions to the experimentally determined target genes of 200 TFs from single-gene perturbation studies in the CREEDS database (knockdown, knockout, or over-expression of a TF, followed by gene expression profiling with microarrays or RNA-seq). x-axis indicates the total number of predicted TF-target gene interactions supported by the single-gene perturbation studies. Arrow indicates the observed count from the TRN model. Histogram indicates an empirical null distribution derived from 10,000 permutations of edges from the TRN model. The observed value was never achieved in 10,000 permutations.

**Figure S3.**
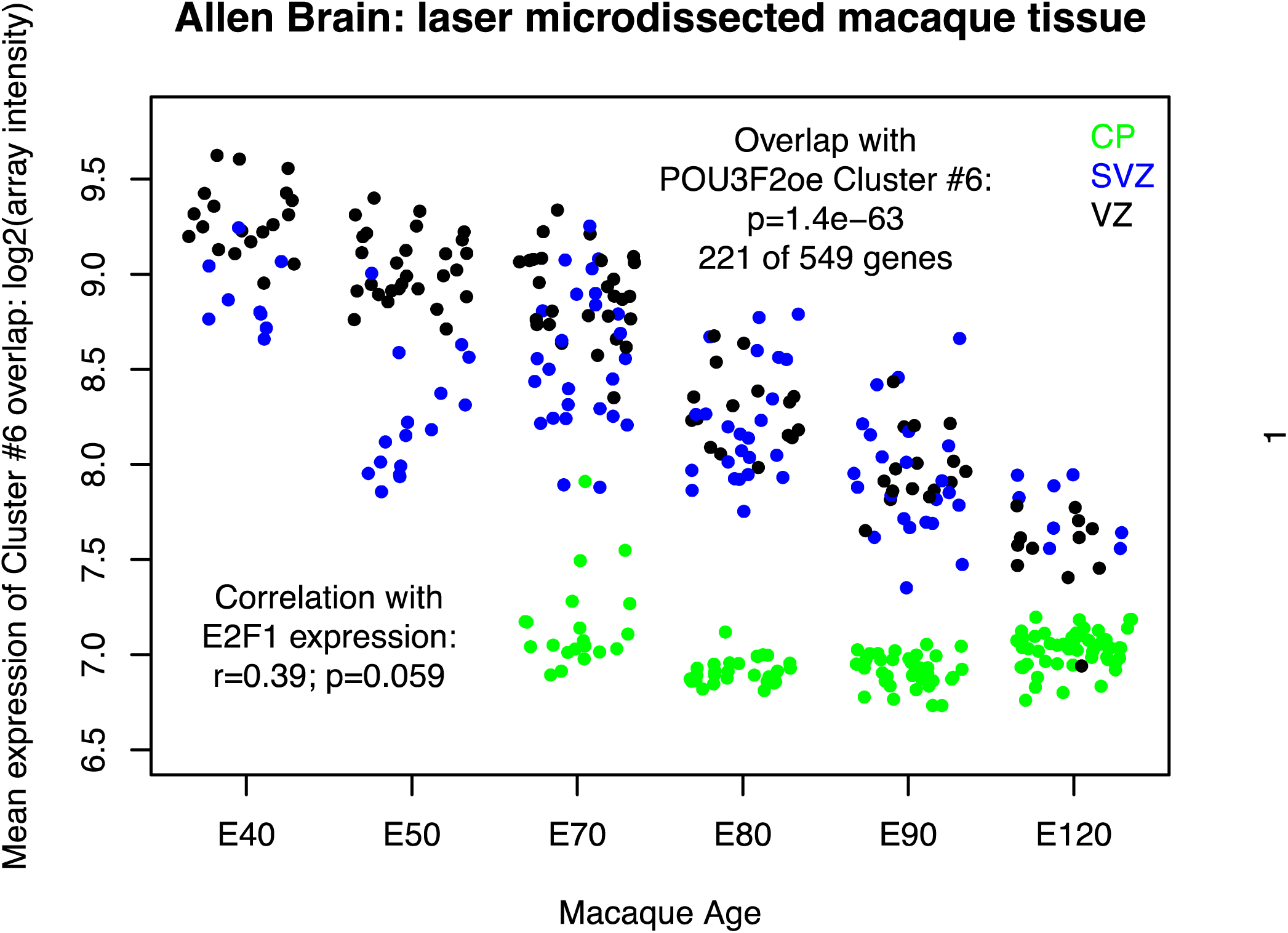
Genetic associations with risk for schizophrenia at the *SREBF1* locus. Region plot was generated with data from Ripke et al. (2014)^14^, using Ricopili (https://data.broadinstitute.org/mpg/ricopili/).

**Figure S4.**
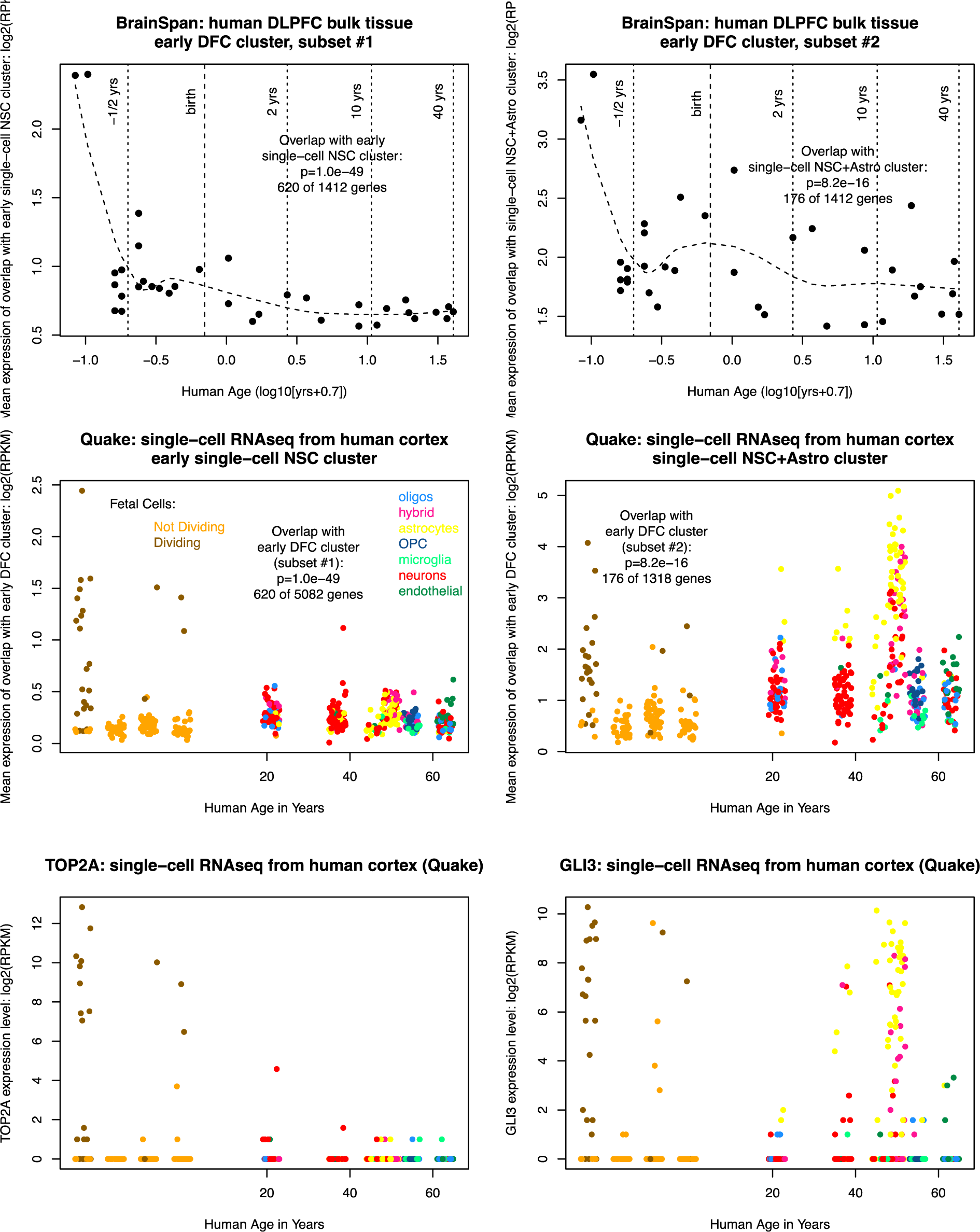
Mean expression of a C6-overlapping cluster in the developing cortex of the non-human primate. We applied k-means clustering to cortical data from the NIH Blueprint Non-Human Primate Brain Atlas^70^ (http://www.blueprintnhpatlas.org), identifying a single gene cluster that overlaps cluster C6 from over-expression of POU3F2 in primary human neural stem cells. Figure S3 shows mean expression levels of the genes in this cluster in laser microdissected non-human primate brain tissue samples. Genes in this cluster had very high expression levels in the VZ and SVZ, and low levels in the developing cortical plate. In contrast to the BrainSpan RNA-seq of developing human brain, the non-human primate dataset captures a larger number of early fetal time points with a gradual decline in the expression of the C6-overlapping genes.

**Figure S5.**
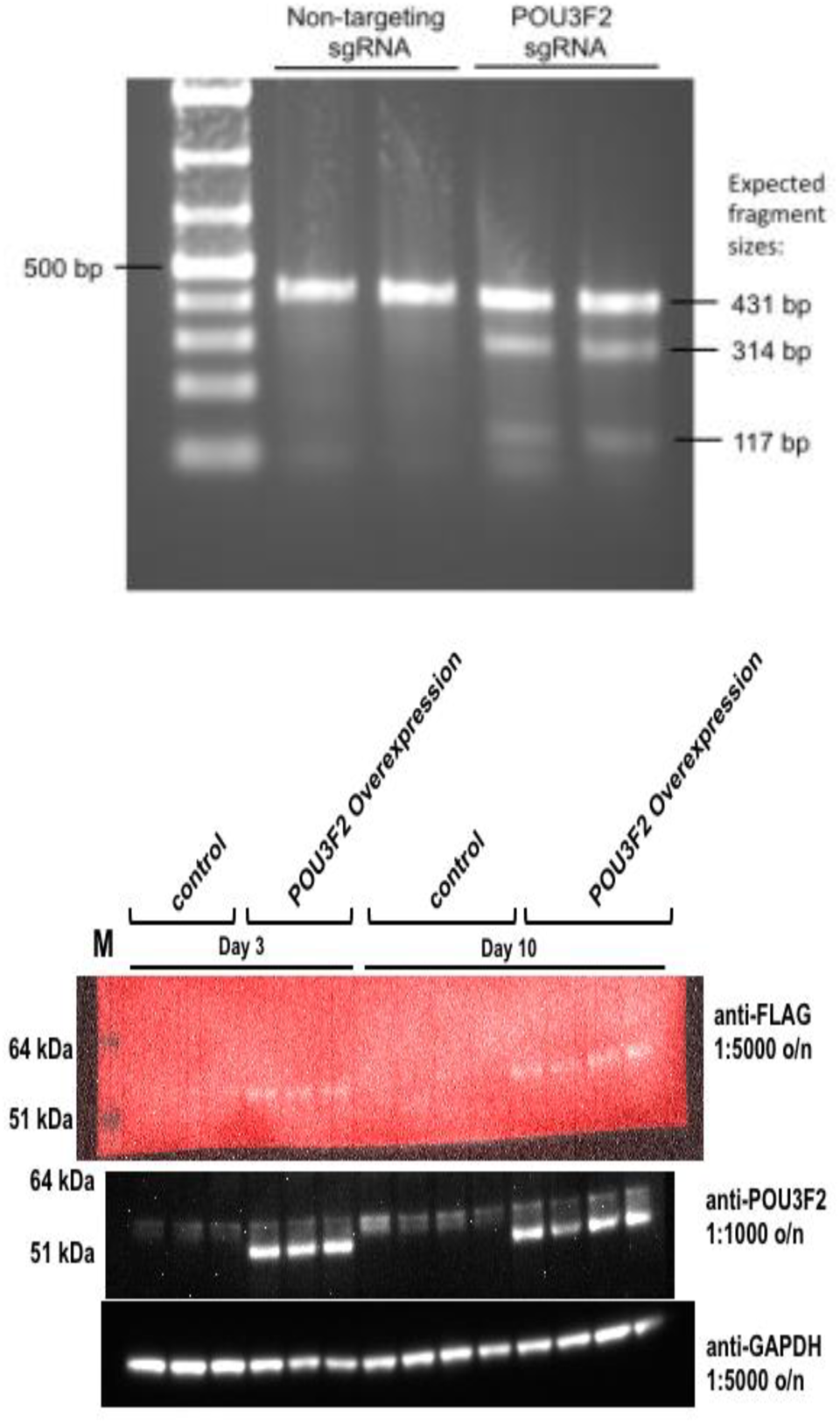
A subset of genes in the C6 cluster are highly expressed in adult astrocytes. Sub-clustering of the C6-overlapping gene clusters in *in vitro* datasets revealed that subsets of the genes in cluster C6 are expressed both in neural stem cells and in adult astrocytes. The preservation of this gene cluster in adult astrocytes may have allowed us to detect stem cell-enriched POU3F2 target genes while studying adult brain tissue.

**Figure S6.**
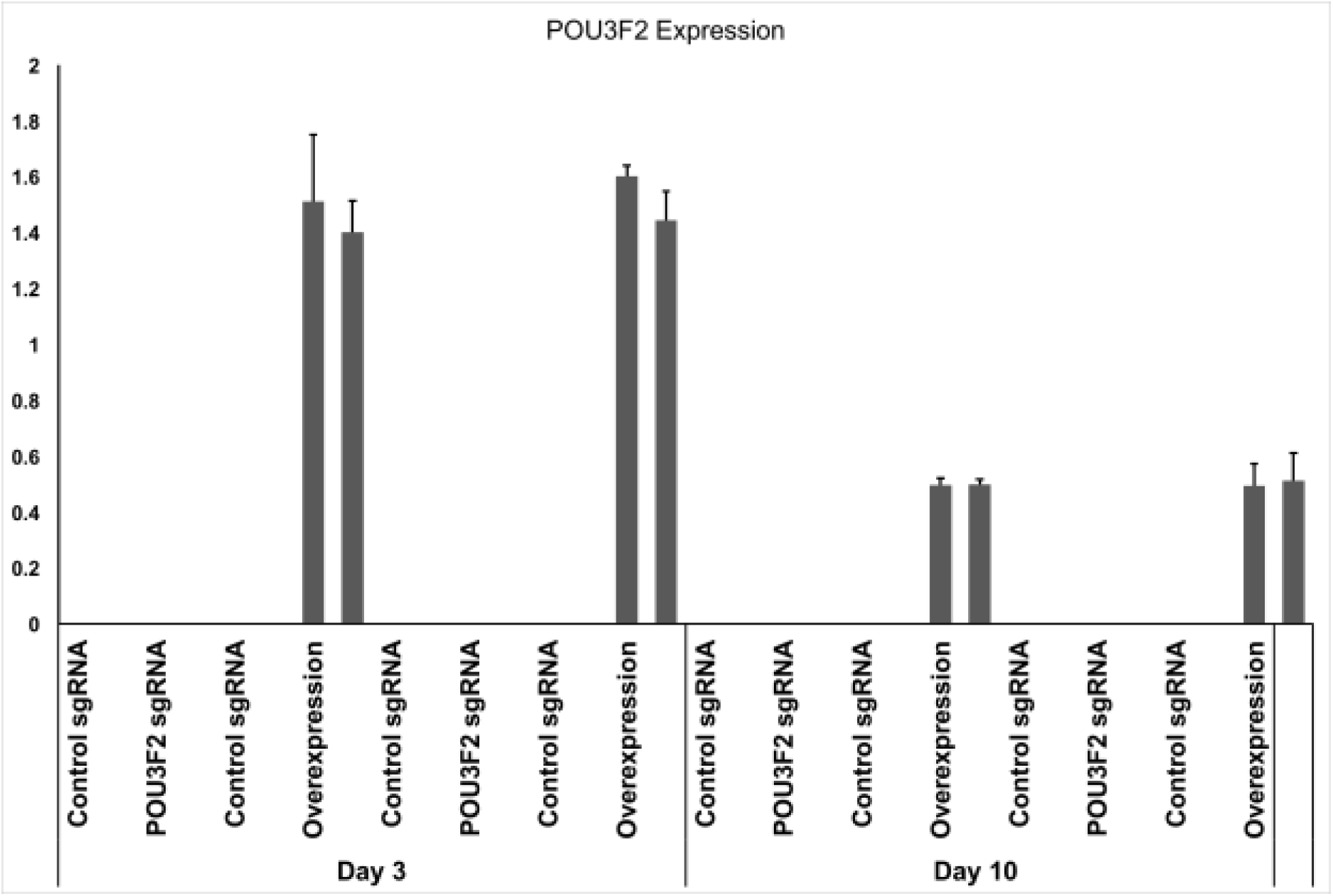
Surveyor Nuclease Assay validation of CRISPR-Cas9 POU3F2 sgRNAs from genomic DNA of transduced human neural stem cells.

Figure S7. Western blot results from overexpression of FLAG-tagged POU3F2 in hNSCs

Figure S8. qPCR validation of POU3F2 overexpression in hNSCs

**Table S1.** Footprinting summary statistics

| Sample ID | Brain Region | mapped reads | DHS regions (min 400 bp) | footprints |
| --- | --- | --- | --- | --- |
| ENCSR000EIJ | cerebellum | 261386073 | 166739 | 51981 |
| ENCSR000EIK | frontal cortex | 192478864 | 53454 | 11834 |
| ENCSR000EIY | frontal cortex | 174008603 | 43132 | 8668 |
| ENCSR000ENA | hippocampus | 543478591 | 180260 | 109701 |
| ENCSR000ENC | cerebellum | 98313605 | 128819 | 21062 |
| ENCSR000ENE | brain microvascular endothelial cell | 97621556 | 140888 | 14582 |
| ENCSR000ENF | brain pericyte | 56208093 | 137903 | 8546 |
| ENCSR000ENL | choroid plexus epithelial cell | 376999221 | 179328 | 138085 |
| ENCSR224IYD | medulla oblongata | 1004733648 | 227548 | 98295 |
| ENCSR318PRQ | middle frontal gyrus | 593844336 | 469949 | 87964 |
| ENCSR475VQD | fetal brain | 486381162 | 218277 | 76210 |
| ENCSR503HIB | cerebellar cortex | 680131240 | 129976 | 88658 |
| ENCSR595CSH | fetal brain | 719039696 | 166037 | 120623 |
| ENCSR706IDL | midbrain | 1021179488 | 300389 | 249300 |
| ENCSR771DAX | globus pallidus | 484128221 | 94788 | 36161 |

**Table S2.**
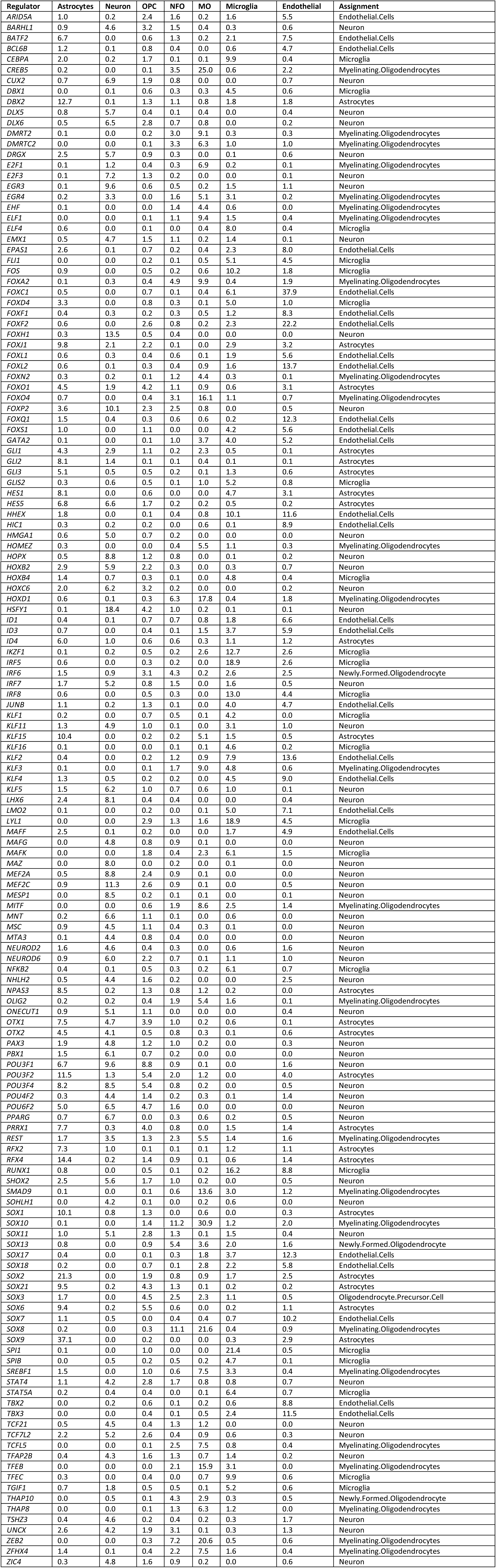
Enrichments of each TF’s target genes for cell type-specific genes.

**Table S3.**
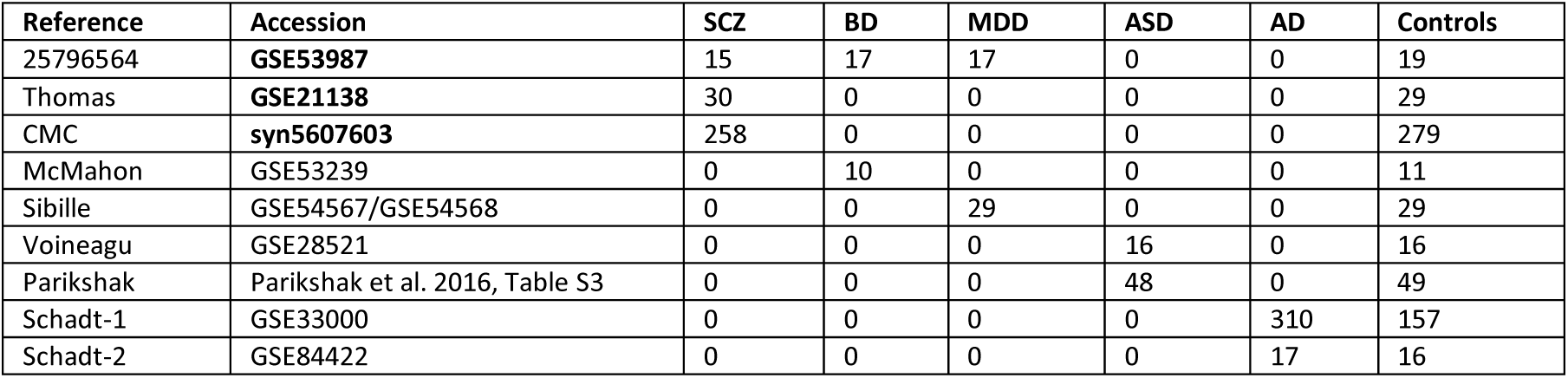
Datasets and accession numbers used for key regulator analysis

**Table S4.** Key regulator TFs for prefrontal cortex gene expression changes in bipolar disorder

| TF | p.up<br>GSE53239 | p.up<br>GSE53987 | meta-p<br>up | q up | p.down<br>GSE53239 | p.down<br>GSE53987 | meta-p<br>down | q down |
| --- | --- | --- | --- | --- | --- | --- | --- | --- |
| *SOX9* | 2.1E-24 | 1.5E-04 | 2.0E-26 | 1.5E-23 | 9.0E-01 | 1.0E+00 | 9.9E-01 | 1.0E+00 |
| *SOX2* | 8.1E-17 | 3.6E-06 | 1.5E-20 | 1.1E-17 | 3.4E-01 | 1.0E+00 | 7.0E-01 | 1.0E+00 |
| *FOXJ1* | 1.6E-15 | 2.8E-07 | 2.3E-20 | 1.7E-17 | 9.6E-01 | 8.9E-01 | 9.9E-01 | 1.0E+00 |
| *FOXO1* | 7.9E-07 | 4.4E-10 | 1.3E-14 | 9.3E-12 | 8.4E-01 | 9.6E-01 | 9.8E-01 | 1.0E+00 |
| *PRRX1* | 5.5E-11 | 8.7E-06 | 1.7E-14 | 1.3E-11 | 2.4E-01 | 9.3E-01 | 5.6E-01 | 1.0E+00 |
| *HMBOX1* | 2.1E-03 | 1.0E-08 | 5.6E-10 | 4.1E-07 | 7.5E-01 | 9.1E-01 | 9.4E-01 | 1.0E+00 |
| *FOXN2* | 2.0E-02 | 7.4E-09 | 3.6E-09 | 2.6E-06 | 9.9E-01 | 9.7E-01 | 1.0E+00 | 1.0E+00 |
| *POU3F4* | 1.5E-06 | 1.4E-03 | 4.6E-08 | 3.3E-05 | 9.1E-01 | 8.0E-01 | 9.6E-01 | 1.0E+00 |
| *POU3F2* | 6.4E-06 | 4.4E-03 | 5.2E-07 | 3.7E-04 | 4.7E-01 | 3.0E-01 | 4.2E-01 | 1.0E+00 |
| *IRF9* | 6.2E-04 | 8.9E-05 | 9.7E-07 | 7.0E-04 | 4.8E-01 | 9.6E-01 | 8.2E-01 | 1.0E+00 |
| *FOXN3* | 2.8E-02 | 4.9E-06 | 2.3E-06 | 1.6E-03 | 8.5E-01 | 9.6E-01 | 9.8E-01 | 1.0E+00 |
| *NPAS3* | 2.5E-05 | 8.7E-03 | 3.6E-06 | 2.6E-03 | 7.9E-01 | 9.2E-01 | 9.6E-01 | 1.0E+00 |
| *RUNX1* | 2.7E-03 | 6.3E-04 | 2.4E-05 | 1.7E-02 | 3.8E-02 | 1.6E-01 | 3.8E-02 | 1.0E+00 |

**Table S5.** Key regulator TFs for prefrontal cortex gene expression changes in schizophrenia

| TF | p.up<br>GSE53987 | p.up<br>GSE21138 | p.up<br>CMC | meta-p<br>up | q up | p.down<br>GSE53987 | p.down<br>GSE21138 | p.down<br>CMC | meta-p<br>down | q down |
| --- | --- | --- | --- | --- | --- | --- | --- | --- | --- | --- |
| *SOX9* | 6.4E-33 | 2.9E-24 | 6.4E-01 | 9.9E-53 | 7.4E-50 | 9.8E-01 | 8.7E-01 | 8.6E-01 | 1.0E+00 | 1.0E+00 |
| *SOX2* | 3.7E-23 | 6.7E-14 | 9.5E-01 | 8.2E-33 | 6.1E-30 | 1.0E+00 | 1.0E+00 | 7.8E-01 | 1.0E+00 | 1.0E+00 |
| *KLF15* | 2.4E-07 | 2.2E-14 | 1.0E+00 | 5.8E-18 | 4.3E-15 | 1.0E+00 | 1.0E+00 | 1.8E-02 | 2.4E-01 | 1.0E+00 |
| *FOXO1* | 1.1E-12 | 1.6E-08 | 4.6E-01 | 8.9E-18 | 6.6E-15 | 1.0E+00 | 2.1E-01 | 1.0E+00 | 8.0E-01 | 1.0E+00 |
| *HES1* | 1.2E-07 | 1.9E-11 | 1.0E+00 | 2.0E-15 | 1.5E-12 | 9.9E-01 | 1.0E+00 | 2.8E-01 | 8.6E-01 | 1.0E+00 |
| *FOXN2* | 3.5E-08 | 3.4E-07 | 6.1E-03 | 5.2E-14 | 3.9E-11 | 8.9E-01 | 9.9E-01 | 1.0E+00 | 1.0E+00 | 1.0E+00 |
| *MAZ* | 1.0E+00 | 1.0E+00 | 9.8E-01 | 1.0E+00 | 1.0E+00 | 1.0E-10 | 8.7E-01 | 1.7E-06 | 1.1E-13 | 8.0E-11 |
| *FOXJ1* | 7.9E-08 | 4.9E-09 | 4.3E-01 | 1.2E-13 | 8.7E-11 | 8.4E-01 | 2.3E-02 | 1.0E+00 | 2.5E-01 | 1.0E+00 |
| *RFX4* | 1.2E-04 | 1.1E-11 | 9.1E-01 | 7.4E-13 | 5.5E-10 | 1.0E+00 | 7.6E-01 | 1.5E-02 | 1.8E-01 | 1.0E+00 |
| *PRRX1* | 6.5E-09 | 2.6E-07 | 1.0E+00 | 1.0E-12 | 7.5E-10 | 8.7E-01 | 9.0E-01 | 9.4E-01 | 1.0E+00 | 1.0E+00 |
| *STAT5B* | 1.9E-05 | 9.5E-04 | 2.2E-07 | 2.3E-12 | 1.7E-09 | 4.4E-01 | 5.9E-01 | 9.9E-01 | 8.4E-01 | 1.0E+00 |
| *ADNP* | 2.9E-05 | 2.0E-04 | 7.4E-07 | 2.5E-12 | 1.8E-09 | 9.9E-01 | 7.2E-01 | 1.0E+00 | 9.9E-01 | 1.0E+00 |
| *ATF2* | 9.4E-01 | 1.0E+00 | 6.1E-02 | 4.5E-01 | 1.0E+00 | 5.4E-07 | 9.6E-09 | 1.0E+00 | 3.0E-12 | 2.2E-09 |
| *IKZF2* | 5.5E-06 | 2.4E-03 | 1.8E-06 | 1.3E-11 | 9.4E-09 | 1.0E+00 | 9.2E-01 | 1.0E+00 | 1.0E+00 | 1.0E+00 |
| *TEAD1* | 4.1E-09 | 2.4E-04 | 1.7E-01 | 7.7E-11 | 5.7E-08 | 2.8E-01 | 4.8E-01 | 8.5E-01 | 6.3E-01 | 1.0E+00 |
| *ZEB1* | 3.6E-06 | 8.9E-07 | 3.6E-01 | 4.7E-10 | 3.4E-07 | 7.4E-01 | 3.2E-01 | 8.2E-01 | 7.7E-01 | 1.0E+00 |
| *NFE2L2* | 7.2E-07 | 3.2E-06 | 5.9E-01 | 5.5E-10 | 4.0E-07 | 9.4E-01 | 8.8E-01 | 8.3E-01 | 9.9E-01 | 1.0E+00 |
| *DBX2* | 7.6E-07 | 8.7E-06 | 2.2E-01 | 5.8E-10 | 4.3E-07 | 9.8E-01 | 5.1E-01 | 1.6E-01 | 5.3E-01 | 1.0E+00 |
| *HMBOX1* | 2.0E-07 | 1.4E-04 | 2.7E-01 | 2.5E-09 | 1.9E-06 | 5.8E-01 | 5.6E-01 | 1.0E+00 | 9.0E-01 | 1.0E+00 |
| *NKX2-1* | 2.0E-05 | 1.5E-05 | 5.1E-02 | 5.3E-09 | 3.8E-06 | 7.9E-01 | 7.8E-01 | 9.5E-01 | 9.8E-01 | 1.0E+00 |
| *ARID4B* | 3.8E-07 | 6.6E-02 | 6.5E-04 | 5.6E-09 | 4.1E-06 | 4.7E-01 | 8.8E-01 | 1.0E+00 | 9.4E-01 | 1.0E+00 |
| *RFX2* | 3.4E-03 | 5.1E-09 | 1.0E+00 | 5.8E-09 | 4.2E-06 | 9.6E-01 | 9.9E-01 | 4.8E-02 | 4.1E-01 | 1.0E+00 |
| *ATRX* | 1.7E-04 | 8.3E-02 | 1.6E-06 | 7.7E-09 | 5.5E-06 | 8.0E-01 | 2.7E-02 | 1.0E+00 | 2.6E-01 | 1.0E+00 |
| *HES6* | 9.9E-01 | 1.0E+00 | 1.0E+00 | 1.0E+00 | 1.0E+00 | 2.3E-03 | 9.8E-01 | 1.1E-08 | 8.1E-09 | 6.0E-06 |
| *ZBTB7A* | 1.0E+00 | 9.7E-01 | 9.9E-01 | 1.0E+00 | 1.0E+00 | 1.5E-03 | 9.9E-01 | 1.8E-08 | 8.7E-09 | 6.4E-06 |
| *SREBF1* | 6.5E-06 | 4.2E-06 | 1.0E+00 | 8.8E-09 | 6.3E-06 | 9.2E-01 | 1.0E+00 | 3.0E-02 | 3.0E-01 | 1.0E+00 |
| *MEIS2* | 3.8E-05 | 1.3E-06 | 8.0E-01 | 1.2E-08 | 8.9E-06 | 7.0E-01 | 2.3E-02 | 7.4E-01 | 1.8E-01 | 1.0E+00 |
| *SOX21* | 8.7E-07 | 2.7E-04 | 2.3E-01 | 1.6E-08 | 1.2E-05 | 9.8E-01 | 7.5E-02 | 9.0E-01 | 4.9E-01 | 1.0E+00 |
| *RUNX1* | 1.3E-03 | 5.2E-08 | 1.0E+00 | 2.1E-08 | 1.5E-05 | 9.0E-01 | 9.6E-01 | 9.3E-01 | 1.0E+00 | 1.0E+00 |
| *NFIC* | 1.0E+00 | 3.9E-01 | 1.0E+00 | 9.3E-01 | 1.0E+00 | 7.5E-03 | 1.3E-01 | 1.2E-07 | 3.2E-08 | 2.4E-05 |
| *LIN54* | 4.6E-04 | 4.6E-01 | 1.6E-06 | 8.6E-08 | 6.2E-05 | 3.6E-01 | 1.2E-01 | 1.0E+00 | 4.0E-01 | 1.0E+00 |
| *TAF1* | 1.3E-04 | 9.8E-05 | 6.3E-02 | 1.9E-07 | 1.4E-04 | 5.6E-01 | 7.3E-01 | 9.9E-01 | 9.4E-01 | 1.0E+00 |
| *TFE3* | 9.9E-01 | 6.7E-01 | 1.0E+00 | 9.9E-01 | 1.0E+00 | 1.2E-03 | 8.2E-01 | 1.0E-06 | 2.2E-07 | 1.7E-04 |
| *NR2F1* | 6.4E-01 | 9.3E-01 | 9.9E-01 | 9.8E-01 | 1.0E+00 | 5.2E-05 | 1.4E-02 | 1.4E-03 | 2.3E-07 | 1.7E-04 |
| *TFAP2C* | 1.0E+00 | 8.2E-01 | 9.8E-01 | 1.0E+00 | 1.0E+00 | 5.1E-03 | 4.9E-01 | 4.0E-07 | 2.4E-07 | 1.8E-04 |
| *NR1H2* | 1.0E+00 | 1.0E+00 | 1.0E+00 | 1.0E+00 | 1.0E+00 | 3.7E-02 | 5.8E-01 | 6.5E-08 | 3.2E-07 | 2.4E-04 |
| *TRPS1* | 1.6E-05 | 1.5E-04 | 9.1E-01 | 4.9E-07 | 3.5E-04 | 9.9E-01 | 2.4E-01 | 5.9E-01 | 6.8E-01 | 1.0E+00 |
| *ID4* | 1.2E-03 | 2.1E-06 | 9.4E-01 | 5.1E-07 | 3.6E-04 | 9.4E-01 | 8.0E-01 | 1.8E-02 | 2.0E-01 | 1.0E+00 |
| *NPAS3* | 1.1E-06 | 4.9E-03 | 8.9E-01 | 1.0E-06 | 7.2E-04 | 9.7E-01 | 9.1E-01 | 9.2E-03 | 1.4E-01 | 1.0E+00 |
| *JUN* | 3.0E-03 | 6.0E-05 | 2.9E-02 | 1.1E-06 | 7.6E-04 | 8.3E-02 | 5.1E-01 | 9.6E-01 | 3.8E-01 | 1.0E+00 |
| *ZHX1* | 1.4E-03 | 2.1E-03 | 2.0E-03 | 1.2E-06 | 8.6E-04 | 8.7E-01 | 2.4E-01 | 9.9E-01 | 7.9E-01 | 1.0E+00 |
| *LYL1* | 6.9E-04 | 1.1E-05 | 1.0E+00 | 1.4E-06 | 1.0E-03 | 9.6E-01 | 9.9E-01 | 1.0E-01 | 6.0E-01 | 1.0E+00 |
| *PATZ1* | 3.5E-03 | 2.4E-06 | 9.1E-01 | 1.5E-06 | 1.0E-03 | 9.7E-01 | 1.0E+00 | 2.0E-01 | 7.8E-01 | 1.0E+00 |
| *FOXO4* | 1.8E-04 | 3.0E-04 | 1.4E-01 | 1.5E-06 | 1.1E-03 | 7.5E-01 | 7.4E-01 | 8.3E-01 | 9.6E-01 | 1.0E+00 |
| *ONECUT1* | 7.3E-03 | 3.7E-01 | 3.5E-06 | 1.8E-06 | 1.3E-03 | 9.7E-01 | 2.8E-01 | 9.9E-01 | 8.5E-01 | 1.0E+00 |
| *IRF9* | 8.5E-05 | 1.4E-04 | 8.3E-01 | 1.8E-06 | 1.3E-03 | 9.8E-01 | 6.0E-01 | 8.7E-01 | 9.7E-01 | 1.0E+00 |
| *MLX* | 1.0E+00 | 1.0E+00 | 9.9E-01 | 1.0E+00 | 1.0E+00 | 8.7E-06 | 8.7E-01 | 1.4E-03 | 2.0E-06 | 1.5E-03 |
| *SMAD9* | 1.0E-05 | 1.3E-03 | 1.0E+00 | 2.3E-06 | 1.7E-03 | 1.0E+00 | 9.7E-01 | 6.3E-02 | 4.7E-01 | 1.0E+00 |
| *NR1H3* | 9.5E-01 | 7.9E-01 | 1.0E+00 | 1.0E+00 | 1.0E+00 | 2.8E-02 | 8.3E-01 | 6.4E-07 | 2.7E-06 | 2.0E-03 |
| *MEF2A* | 7.4E-01 | 1.5E-01 | 1.8E-06 | 2.7E-05 | 1.9E-02 | 3.3E-06 | 6.3E-03 | 1.0E+00 | 3.6E-06 | 2.7E-03 |
| *ARID1A* | 3.5E-05 | 3.3E-02 | 1.8E-02 | 3.7E-06 | 2.6E-03 | 9.2E-01 | 7.2E-01 | 9.0E-01 | 9.8E-01 | 1.0E+00 |
| *TSHZ3* | 1.0E+00 | 9.9E-01 | 3.3E-01 | 9.0E-01 | 1.0E+00 | 1.7E-04 | 3.9E-02 | 3.5E-03 | 4.0E-06 | 2.9E-03 |
| *KLF9* | 9.9E-01 | 1.3E-01 | 8.7E-01 | 6.3E-01 | 1.0E+00 | 3.7E-03 | 7.9E-01 | 9.4E-06 | 4.6E-06 | 3.4E-03 |
| *POU3F4* | 3.0E-05 | 5.9E-03 | 1.6E-01 | 4.8E-06 | 3.3E-03 | 8.5E-01 | 4.9E-01 | 8.6E-01 | 9.1E-01 | 1.0E+00 |
| *KLF13* | 1.0E+00 | 3.6E-01 | 9.2E-01 | 9.0E-01 | 1.0E+00 | 1.2E-02 | 9.5E-01 | 2.7E-06 | 5.2E-06 | 3.8E-03 |
| *NCOA1* | 1.0E+00 | 6.8E-01 | 1.0E+00 | 9.9E-01 | 1.0E+00 | 5.1E-05 | 5.7E-01 | 1.5E-03 | 6.9E-06 | 5.0E-03 |
| *ARID4A* | 1.1E-02 | 4.2E-01 | 9.8E-06 | 7.3E-06 | 5.1E-03 | 7.3E-01 | 9.8E-01 | 9.9E-01 | 9.9E-01 | 1.0E+00 |
| *PPARA* | 7.7E-05 | 6.1E-04 | 9.9E-01 | 7.5E-06 | 5.3E-03 | 9.9E-01 | 9.1E-01 | 1.0E+00 | 1.0E+00 | 1.0E+00 |
| *FOXN3* | 1.7E-02 | 3.1E-02 | 1.1E-04 | 9.3E-06 | 6.5E-03 | 2.2E-01 | 4.1E-01 | 1.0E+00 | 5.7E-01 | 1.0E+00 |
| *POU3F2* | 2.4E-03 | 7.7E-04 | 3.4E-02 | 9.7E-06 | 6.8E-03 | 7.4E-01 | 4.5E-01 | 9.7E-01 | 8.9E-01 | 1.0E+00 |
| *NR2F6* | 1.0E+00 | 9.8E-01 | 1.0E+00 | 1.0E+00 | 1.0E+00 | 5.7E-03 | 9.6E-01 | 1.5E-05 | 1.2E-05 | 8.8E-03 |
| *SP3* | 1.5E-02 | 8.3E-03 | 8.0E-04 | 1.5E-05 | 1.0E-02 | 5.5E-01 | 7.5E-01 | 1.8E-02 | 1.3E-01 | 1.0E+00 |
| *FOXC1* | 3.5E-02 | 3.5E-06 | 8.8E-01 | 1.6E-05 | 1.1E-02 | 1.6E-01 | 9.8E-01 | 6.0E-01 | 5.8E-01 | 1.0E+00 |
| *FOXO3* | 1.0E-04 | 2.0E-01 | 6.9E-03 | 2.0E-05 | 1.4E-02 | 1.4E-01 | 2.1E-01 | 1.0E+00 | 3.1E-01 | 1.0E+00 |
| *JDP2* | 9.6E-01 | 9.1E-01 | 1.2E-01 | 6.0E-01 | 1.0E+00 | 3.5E-04 | 1.2E-02 | 4.7E-02 | 2.7E-05 | 1.9E-02 |
| *SMAD1* | 2.6E-05 | 2.5E-02 | 3.6E-01 | 3.1E-05 | 2.1E-02 | 9.5E-01 | 8.6E-01 | 3.6E-01 | 8.7E-01 | 1.0E+00 |
| *FOXH1* | 9.1E-01 | 9.8E-01 | 1.2E-01 | 6.2E-01 | 1.0E+00 | 6.9E-04 | 4.7E-04 | 8.6E-01 | 3.6E-05 | 2.6E-02 |
| *ESRRG* | 9.8E-01 | 1.0E+00 | 7.3E-01 | 1.0E+00 | 1.0E+00 | 5.9E-02 | 1.9E-03 | 3.3E-03 | 4.6E-05 | 3.3E-02 |
| *BPTF* | 3.6E-04 | 1.4E-02 | 7.8E-02 | 4.7E-05 | 3.2E-02 | 9.5E-01 | 9.5E-01 | 9.9E-01 | 1.0E+00 | 1.0E+00 |

**Table S6.**
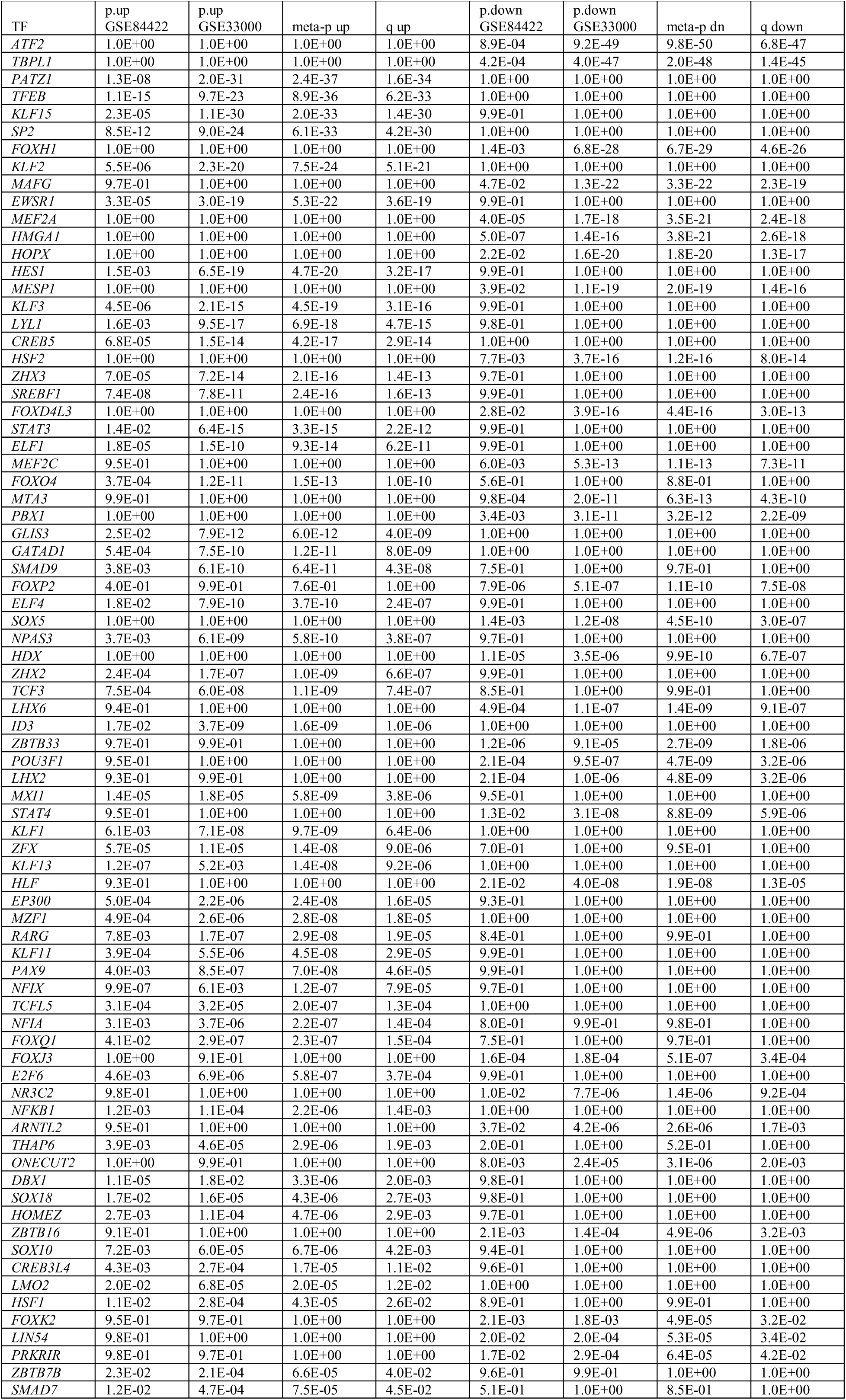
Key regulator TFs for prefrontal cortex gene expression changes in autism spectrum disorder

**Table S7.**
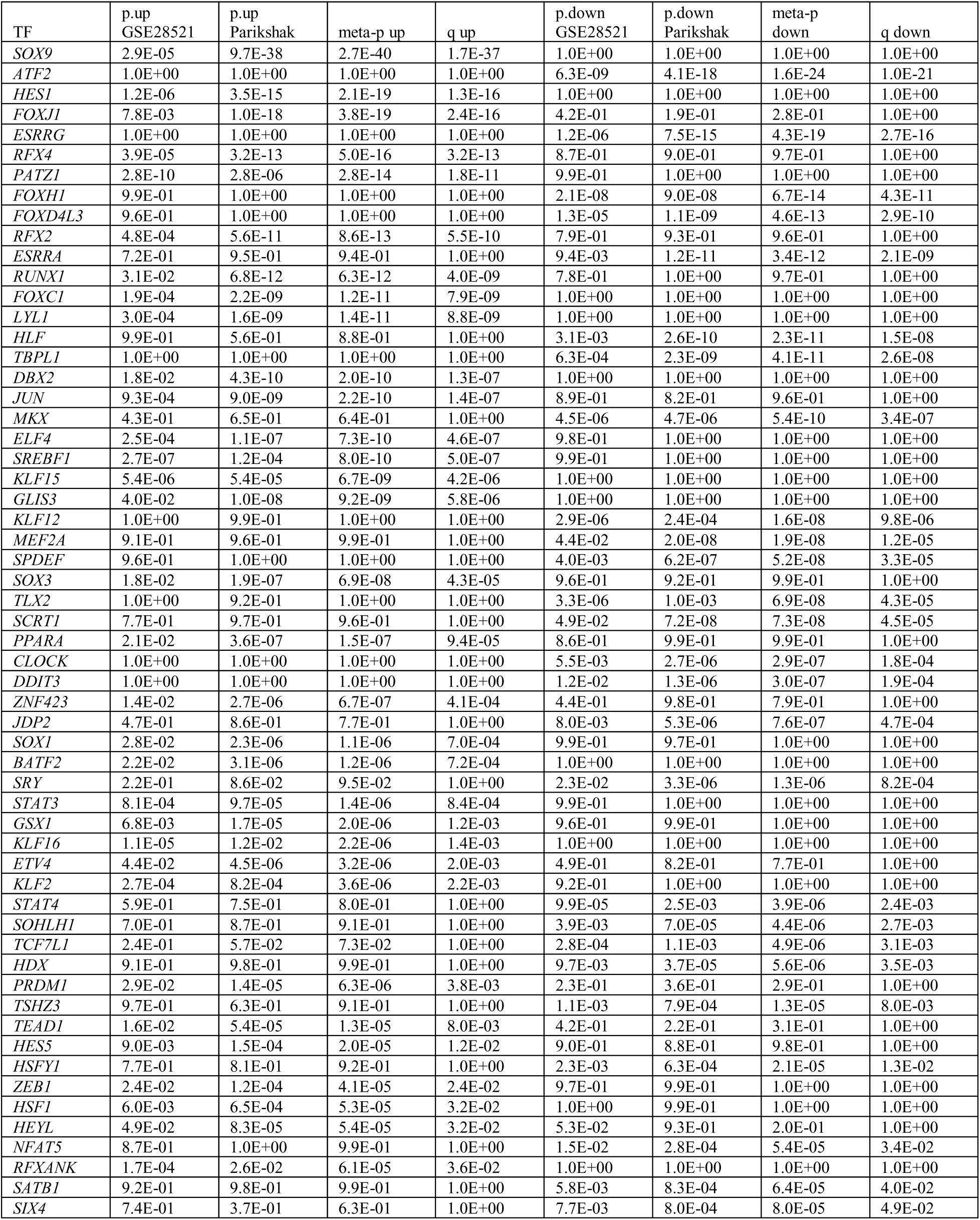
Key regulator TFs for prefrontal cortex gene expression changes in Alzheimer’s disease

**Table S8.** Key regulator TFs at GWAS risk loci

| PubMed ID | First Author | Disease Trait | SNP | P-value | TRN Key<br>Regulator<br>Analysis | TF |
| --- | --- | --- | --- | --- | --- | --- |
| 23562540 | Cruchaga C | Alzheimer's disease biomarkers | rs514716 | 3.00E-09 | AD | GLIS3 |
| 24618891 | Muhleisen | Bipolar disorder | rs12202969 | 1.00E-08 | SCZ,BD | POU3F2 |
| 25056061 | Ripke S | Schizophrenia | rs8082590 | 2.00E-08 | SCZ,AD,ASD | SREBF1 |
| 24162737 | European<br>Alzheimer's<br>Disease Initiative<br>(EADI) | Alzheimer's disease (late onset) | rs190982 | 3.00E-08 | AD | MEF2C |
| 27329760 | Hou L | Bipolar disorder | rs1487441 | 3.00E-08 | SCZ,BD | POU3F2 |
| 26198764 | Goes FS | Schizophrenia | rs9398171 | 4.00E-08 | SCZ | FOXO3 |
| 26198764 | Goes FS | Schizophrenia | rs12883788 | 3.00E-07 | SCZ,BD,AD | NPAS3 |
| 26830138 | Herold C | Alzheimer disease and age of<br>onset | rs3931397 | 6.00E-07 | AD | NR3C2 |
| 20889312 | Wang KS | Bipolar disorder and<br>schizophrenia | rs11880706 | 2.00E-06 | SCZ,ASD | RFX2 |
| 26821981 | Koga AT | antipsychotic drug dosage in<br>schizophrenia or<br>schizoaffective disorder | rs75905933 | 4.00E-06 | SCZ,AD | KLF13 |
| 20713499 | Huang J | Schizophrenia, bipolar disorder<br>and depression (combined) | rs4982029 | 4.00E-06 | SCZ,BD,AD | NPAS3 |
| 18711365 | Ferreira MA | Bipolar disorder | rs8015959 | 5.00E-06 | SCZ,BD,AD | NPAS3 |
| 27770636 | Mez J | Late-onset Alzheimer's disease | rs148003968 | 5.00E-06 | AD | TFEB |
| 23453885 | Smoller JW | Autism spectrum disorder,<br>attention deficit-hyperactivity<br>disorder, bipolar disorder,<br>major depressive disorder, and<br>schizophrenia (combined) | rs7565792 | 7.00E-06 | BD,SCZ | FOXN2 |
| 19023125 | Potkin SG | Brain imaging in schizophrenia<br>(interaction) | rs9491640 | 9.00E-06 | SCZ,BD | POU3F2 |

**Table S9.**
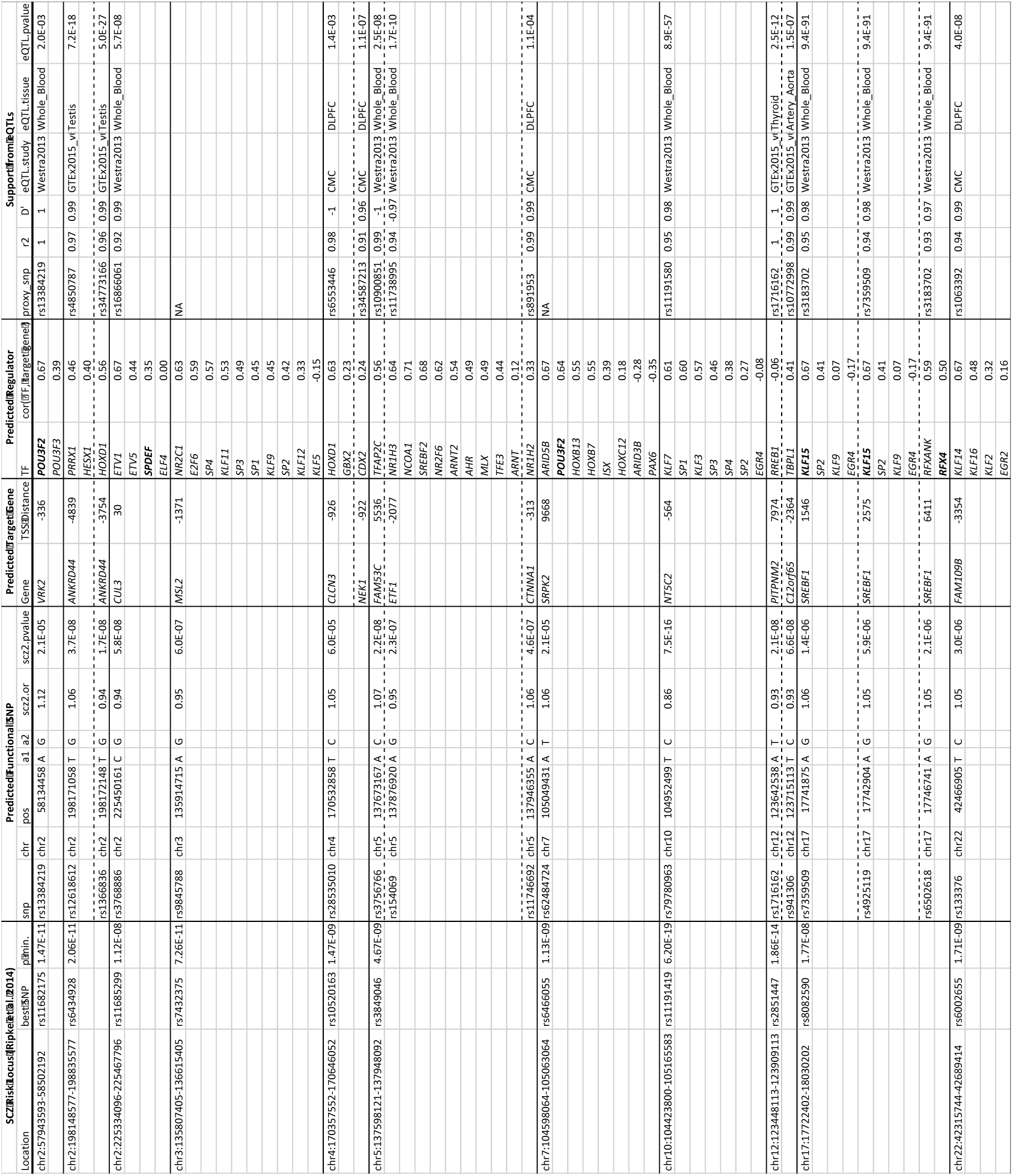
TFBS-disrupting, SCZ-associated SNPs

**Table S10.**
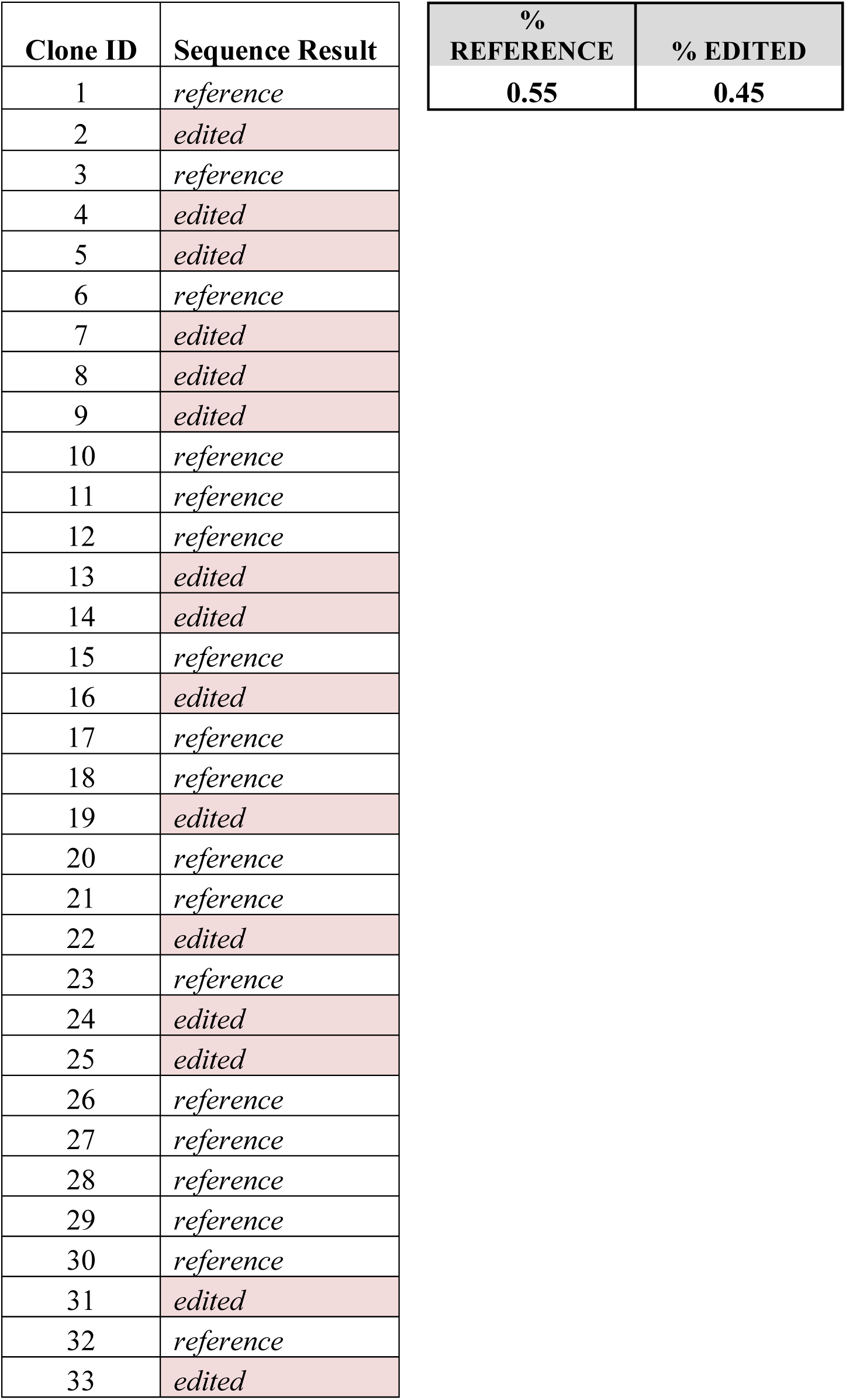
TOPO cloning analysis of POU3F2 sgRNA editing at target site.

## URLs

The following supplementary datasets are available online: http://amentlab.igs.umaryland.edu/psych-trn-pfc2016/

Supplementary Dataset 1: DNase I footprints from brain regions and cell types

Supplementary Dataset 2: Predicted TF-target gene interactions

Supplementary Dataset 3. Variants in TFBSs

Supplementary Dataset 4. POU3F2 clustering results

